# SSnet: A Deep Learning Approach for Protein-Ligand Interaction Prediction

**DOI:** 10.1101/2019.12.20.884841

**Authors:** Niraj Verma, Xingming Qu, Francesco Trozzi, Mohamed Elsaied, Nischal Karki, Yunwen Tao, Brian Zoltowski, Eric C. Larson, Elfi Kraka

## Abstract

Computational prediction of Protein-Ligand Interaction (PLI) is an important step in the modern drug discovery pipeline as it mitigates the cost, time, and resources required to screen novel therapeutics. Deep Neural Networks (DNN) have recently shown excellent performance in PLI prediction. However, the performance is highly dependent on protein and ligand features utilized for the DNN model. Moreover, in current models, the deciphering of how protein features determine the underlying principles that govern PLI is not trivial. In this work, we developed a DNN framework named SSnet that utilizes secondary structure information of proteins extracted as the curvature and torsion of the protein backbone to predict PLI. We demonstrate the performance of SSnet by comparing against a variety of currently popular machine and non-machine learning models using various metrics. We visualize the intermediate layers of SSnet to show a potential latent space for proteins, in particular to extract structural elements in a protein that the model finds influential for ligand binding, which is one of the key features of SSnet. We observed in our study that SSnet learns information about locations in a protein where a ligand can bind including binding sites, allosteric sites and cryptic sites, regardless of the conformation used. We further observed that SSnet is not biased to any specific molecular interaction and extracts the protein fold information critical for PLI prediction. Our work forms an important gateway to the general exploration of secondary structure based deep learning, which is not just confined to protein-ligand interactions, and as such will have a large impact on protein research while being readily accessible for *de novo* drug designers as a standalone package.

## Introduction

Diverse biological processes are dictated by ligand induced conformational changes in target proteins. Modern medicine has harnessed the ability to control protein structure and function through the introduction of small molecules as therapeutic interventions to diseases. Despite the importance of Protein-Ligand Interactions (PLI) in medicine and biology and keen insight into the multitude of factors governing ligand recognition, including hydrogen bonding, ^1,2^ *π*-interactions,^3^ and hydrophobicity,^4^ the development of robust predictive PLI models and validation in drug discovery remains challenging.

Reliance on experimental methods to identify and confirm PLIs is time-consuming and expensive. In contrast, computational methods can save time and resources by filtering large compound libraries to identify smaller subsets of ligands that are likely to bind to the protein of interest. In this manner, reliable PLI predictive algorithms can significantly accelerate the discovery of new treatments, eliminate toxic drug candidates, and efficiently guide medicinal chemistry.^5^ Currently, Virtual Screening (VS) is commonly used in academia and industry as a predictive method of determining PLI. Broadly, VS can be divided into two major categories: Ligand Based Virtual Screening (LBVS) and Structure Based Virtual Screening (SBVS).^6^ LBVS applies sets of known ligands to a target of interest and is therefore limited in its ability to find novel chemotypes. In contrast, SBVS uses the 3D structure of a given target to screen libraries, thereby improving its utility in identifying novel therapeutics. ^7^ Over the last few decades, many classical techniques such as force field, empirical, and knowledge based^5^ PLI predictions have been developed, with limited success. Often these methods show low performance and in some cases even discrepancies when compared with experimental bioactivities.^8^ Even successful methods are often limited by a requirement of high resolution protein structures with detailed information about the binding pocket.^9^

The advent of Machine Learning (ML) and Deep Learning (DL) approaches have created a path towards solving previously challenging unsolved problems in biology and chemistry.^10–17^ Various reviews summarize the application of ML/DL in drug design and discovery.^18–22^ Machine learning based PLI prediction has been developed from a chemogenomic perspective^23^ that considers interactions in a unified framework from chemical space and proteomic space. Some notable examples are: Jacob and Vert ^24^ used tensor-product based features and applied Support Vector Machines (SVM); Yamanishi et al. ^25^ minimized Euclidean distances over common features derived by mapping ligands and proteins; Wallach et al. ^26^ used a 3D grid for proteins along with 3D convolutional networks; Tsubaki et al. ^27^ used a combination of convolutional network for proteins and graph network for ligands; Li et al. ^28^ used Bayesian additive regression trees to predict PLI; and lastly Lee et al. ^29^ applied deep learning with convolution neural networks on protein sequences. While these methods provide novel insights for PLI, they do not provide a solid framework for direct application in drug discovery.

End-to-end learning, a powerful ML/DL technique, has gained interest in recent years since once the model is trained the users are only required to provide standard protein and ligand descriptors as input.^30^ The end-to-end learning technique involves i) embedding inputs to lower dimensions, ii) formulating various neural networks depending on the data available, and iii) using backpropagation over the whole architecture to minimize loss and update weights. An example of an end-to-end learning model that has achieved high level of accuracy in PLIs prediction is GNN-CNN.^27^ Tsubaki et al. ^27^ were able to achieve a remarkable accuracy with only primary sequence information and without any structural insight. However, PLI is highly dependent on the structural assembly of the protein.^1–4^ Since predicting structure of protein from the primary sequence is still an unsolved problem, the ability of the ML/DL to understand structural elements and predict PLI with respect to the ensemble is limited. However, current protein structure-based ML/DL methods for PLI predictions achieve low accuracy as they suffer from three major limitations: i) absence of high resolution structures of protein-ligand pairs for training, ii) the 3D grid for the target can form a huge and sparse matrix, which hinder ML/DL models to learn and predict PLI, and iii) techniques are sensitive to the method employed to represent ligand structure, diverse methods have been reported^31–34^ and selection of the optimal ligand representation can be challenging.

Strategies to overcome these limitations have largely focused on developing new methodologies to reduce target and compound structure space to 1D representations, thereby providing a dense framework for ML/DL to operate on a small number of input features. Reduction of 3D protein structure information into 1D allows the machine to efficiently learn 3D space features of the protein secondary structure, which are required for ligand interaction. This information, when combined with the way a convolution network considers the input space, makes the model unbiased towards the protein conformation, thereby not being limited by the existence of high-resolution protein-ligand structures adequate for PLI prediction. This feature can solve a major drawback for most virtual screening methods since only a limited portion of the proteins’ conformational space can be crystalized.

Herein, we outline a new machine learning based algorithm termed SSnet for the prediction of PLIs. SSnet uses a 1D representation based on the curvature and torsion of the protein backbone. Mathematically, curvature and torsion are sufficient to reasonably represent the 3D structure of a protein^35^ and therefore contain compact information about its function and fold. Further, curvature and torsion are sensitive to slight changes in the secondary structure, which are a consequence of all atom interactions, including side-chains. These characteristics position SSnet to outperform existing methods of PLI prediction to rapidly curate large molecular libraries to identify a subset of likely high-affinity interactors. As outlined below, corresponding analyses are carried out to show the robustness and versatility of our new model. We analyzed the model using the Grad-CAM to visualize heatmaps of the activation from the neural network that maximally excite the input features.^36^ The input features can then be used to highlight on the protein 3D structure the residues that maximally influenced the predicted score.

In the methods and rationale section, we demonstrate how the secondary structure of proteins is used in the ML/DL framework. We discuss the representation of ligands following the introduction of SSnet model, possible evaluation criteria, its merits and demerits. We discuss the datasets used in this work to validate and train SSnet. In the results section, we first select the ligand descriptor to be used in SSnet. We validate SSnet trained on two models named: SSnet:DUD-E, a model trained on DUD-E, ^37^ and SSnet:BDB, a model trained on BDB^38^ for general application. The applicability as a VS tool is demonstrated by using enrichment factor. We further show the applicability of SSnet for a virtual screening task through its high AUCROC and EF score while maintaining a lack of conformational bias allowing it to find ligands for cryptic proteins and visualize important residues considered by SSnet. In the discussion section we outline some key conclusions observed from SSnet and it’s limitations. The conclusion section outlines a way to incorporate SSnet into the drug-design workflow as well as provide a future perspective.

## Methods and Rationale

### Representation of Proteins

Protein structures exhibit a large conformational variety. Many automated and manual sorting databases like SCOP,^39^ CATH,^40^ DALI,^41^ and programs like DSSP,^42^ STRIDE,^43^ DEFINE,^44^ and KAKSI^45^ etc. have provided protein classifications based on the secondary structure. However, these classifications often conflict with each other.^46^ A more promising approach to determine protein fold based on secondary structure has been introduced by Ranganathan et al. ^35^ and Guo and Cremer ^47^ coined Automated Protein Structure Analysis (APSA). The original idea behind APSA is based on Unified Reaction Valley Approach (URVA) developed by Kraka et al. ^48^, where a reaction path is characterized by its arc length, curvature, and torsion. Inspired by the representation of features in APSA, we explored if and how we can utilize a similar secondary structure characterization in ML/DL approaches for PLI prediction.

A protein can be represented by the *α* carbons (CA atoms) of the backbone as it defines a unique and recognizable structure, especially for protein categorization.^49^ In fact, a significant amount of information about the protein is embedded in the secondary structure elements such as helices, *β* sheets, hairpins, coils, and turns etc. Therefore, utilization of these parameters should retain adequate information to train a ML/DL approach.

The secondary structure information can be retrieved by a smooth curve generated by a cubic spline fit of the CA atoms. Figure 1a shows the arc length *s*, scalar curvature *κ*, and scalar torsion *τ* which define the 3D curve **r**(*s*). The scalar curvature *κ* is expressed as a function of arc length *s*

**Figure 1:**
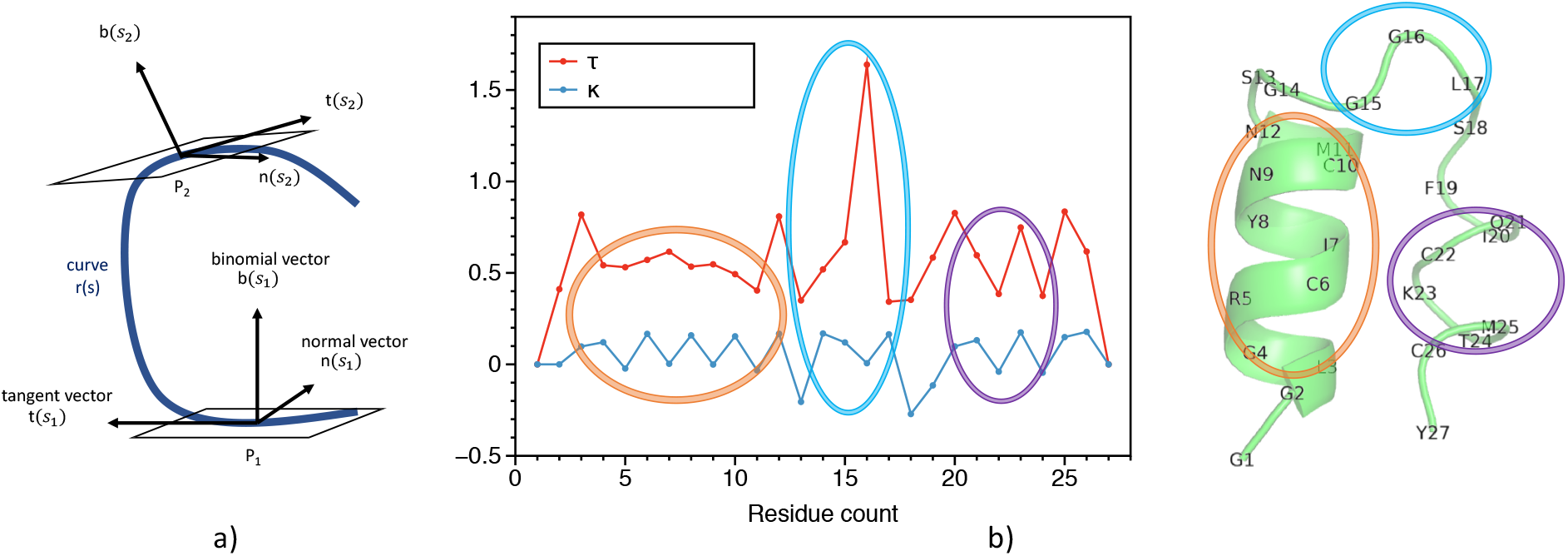
a) The tangent vector **t**, normal vector **n** and the binormal vector **b** of a Frenet frame at points P_1_ and P_2_ respectively for a curve **r**(*s*). b) Representation of protein backbone in terms of scalar curvature *κ* and torsion *τ* respectively. The ideal helix, turn and non-ideal helix is shown in orange, cyan and magenta respectively. The curvature and torsion pattern captures the secondary structure of the protein.

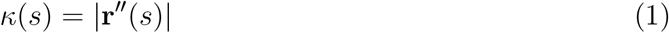

and the scalar torsion

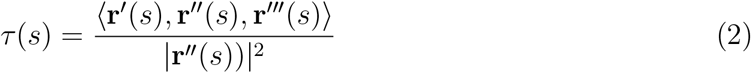

where | ·| is the norm and ⟨·⟩ is the vector triple product. A protein can then be represented by considering the curvature and torsion at the anchor points (locations of CA) forming a 1D vector with twice the the number of amino acids.^35,47^ We limit the number of protein chains to 6 and the number of amino acids per chain as 1500 to have consistent input size. Thus the input size is (9000, 2) i.e. 6 * 1500 for curvature and the same for torsion. The databases used for compiling PLI data for training and testing the models mostly contained 6 chains or lower and 1500 amino acids or lower, therefore 6 chains and 1500 residues already encompass a large amount of proteomic space that might influence ligand binding. Furthermore, DNN can be optimized by having the most dense training and testing dataset and thus 6 chains and 1500 amino acids were used to include the largest amount of data while ensuring that the input is mostly dense.

Figure 1b shows the decomposition of a protein found in Conus villepinii (PDB ID - 6EFE) into scalar curvature *κ* and torsion *τ*, respectively. The residues 5 through 10 show a near ideal *α* helix type secondary structure, which is represented as an oscillation of *τ* with smooth *κ*. Similarly, the turn (residues 15 to 17) and a non-ideal *α* helix (residues 20 to 25) are captured in the decomposition plot via unique patterns. Because the curvature and torsion information of the secondary structure of proteins are encoded as patterns, ML techniques may be powerful tools to predict PLI through efficiently learned representations of these patterns. More specifically, we hypothesize that, using convolution, varying sized filters may be excellent pattern matching methods for discerning structure from these decomposition plots. More analysis on protein representation is provided in the subsection SSnet model.

### Representation of Ligands

A molecule can be represented by the SMILES string, which represents its various bonds and orientations. However, the SMILES string encodes dense information making it difficult for an algorithm to decipher and learn chemical properties. A number of alternative representations for ligands have been proposed that model varying aspects of the ligand in a more machine readable format. The hope is that ML algorithms can more effectively use these representations for prediction. Since ligand representation is an ongoing research topic, we consider four different methods: CLP,^31^ GNN,^34^ Avalon,^32^ and ECF.^33^ CLP was generated by the code provided by Gómez-Bombarelli et al. ^31^; Avalon and ECF were generated from RDKit^50^; and GNN was implemented as proposed by Tsubaki et al. ^27^ where we replace the first dense network in Figure 2 by a graph neural network.

**Figure 2:**
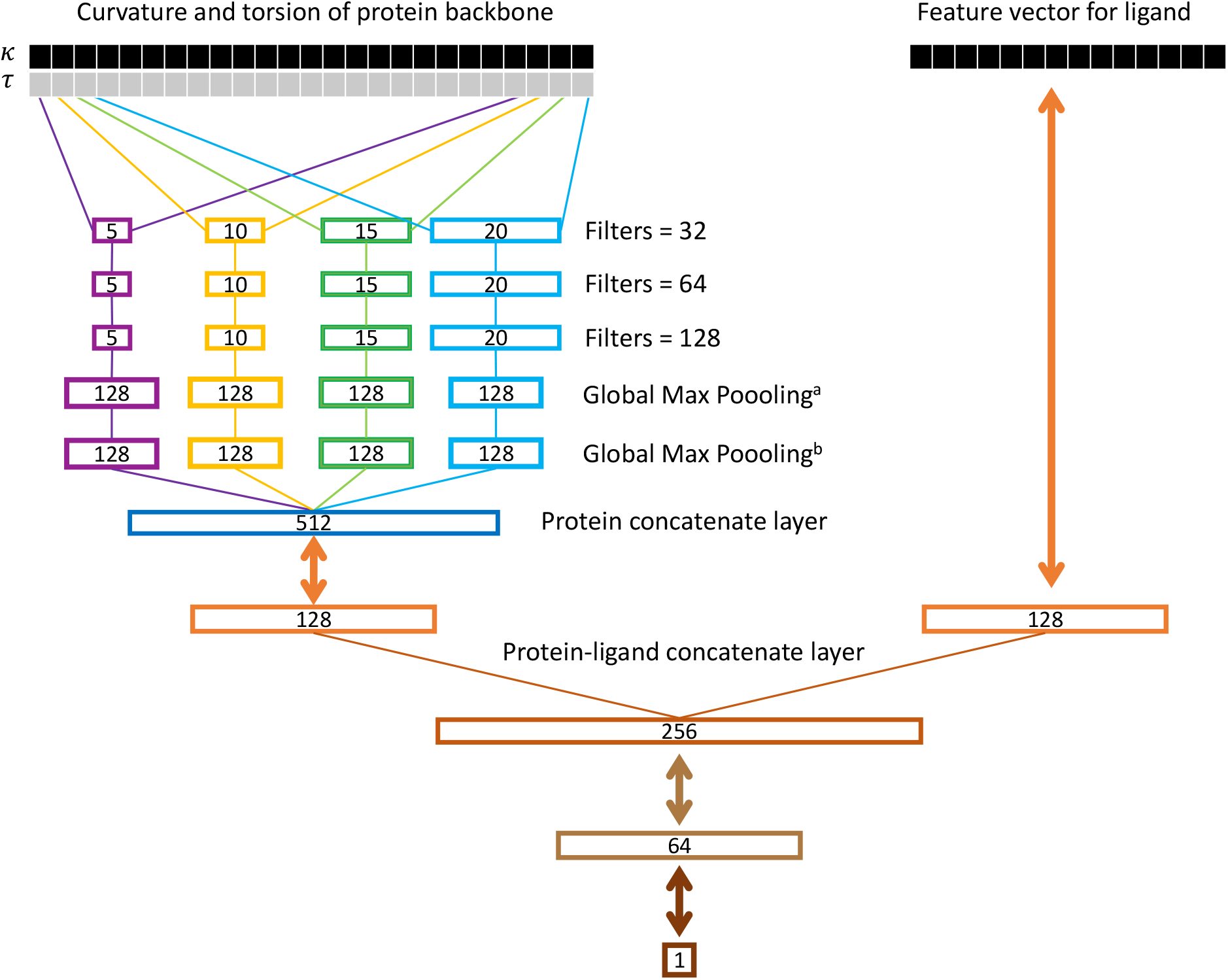
SSnet model. The curvature and torsion pattern of a protein backbone is fed through multiple convolution networks with varying window sizes as branch convolution. Each branch further goes through more convolution with same window size (red, orange, green and light blue boxes). A global max pooling layer is implemented to get the protein vector. The ligand vector is directly fed to the network. Each double array line implies a fully connected dense layer. The number inside a box represents the dimension of the corresponding vector. In the case of GNN, the ligand vector is replaced by a graph neural network as implemented by Tsubaki et al. ^27^

### SSnet model

Figure 2 shows the SSnet model developed in this work. Here, we provide a general overview of the network, more details about its specific design operation are given in the later part of this section. As denoted in the left upper branch of Figure 2, after conversion into the Frenet-Serret frame and the calculation of curvature *κ* and torsion *τ, κ* and *τ* data (i.e decomposition data) is fed into the neural network. We denote this input as a 2D matrix (1D vector with curvature and torsion reshaped to contain curvature in one row and torsion in the other), **X**^(0)^, where each column represents a unique residue and the rows corresponding the curvature and torsion. The first layer is a branch convolution with varying window sizes. That is, each branch is a convolution with a filter of differing length. We perform this operation so that patterns of varying lengths in the decomposition plot can be recognized by the neural network. Each branch is then fed to more convolutions of same window size. This allows the network to recognize more intricate patterns in **X**^(0)^ that might be more difficult to recognize with a single convolution. The output of these convolutional branches are concatenated, pooled over the length of the sequence, and fed to a fully connected dense layer. The rightmost upper branch of Figure 2 shows a ligand vector which is generated and fed to a fully connected dense layer. The output of this layer is typically referred to as an embedding. Intuitively, this embedding is a reduced dimensionality representation of the protein and ligand. The outputs of the protein embedding and the ligand embedding are then concatenated and fed to further dense layers to predict the PLI.

The convolutional network in this research uses filter functions over the protein vector **X**^(0)^. To define the convolution operation more intuitively, we define a reshaping operation as follows:

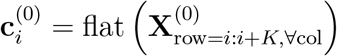

where the flattening operation reshapes the row of **X**^(0)^ from indices *i* to *i*+*K* to be a column vector 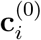. This process is also referred to as vectorization. The size of the filter will then be of length *K*. We define the convolution operation as:

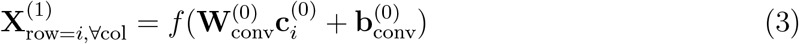

where *f* is a function known as the rectified linear unit (ReLU), 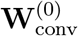 is the weight matrix and 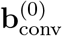 is the bias vector. This operation fills in the columns of the output of the convolution, 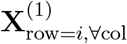 (also called the activation or feature map). Each row of 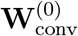 is considered as a different filter and each row of **X**^(1)^ is the convolutional output of each of these filters.

These convolutions can be repeated such that the *n*^*th*^ activation is computed as:

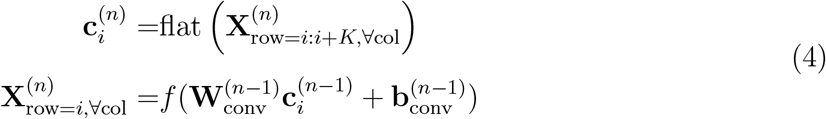

We in our SSnet model use four different branches with filter sizes of *κ* = 5, 10, 15 and 30. The final convolutional activations for layer *N* can be referred to as 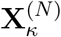 where *κ* denotes the branch. The activation 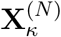 is often referred to as the latent space because it denotes the latent features of the input sequence. The number of columns in 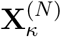 is dependent upon the size of the input sequence. To collapse this unknown size matrix into a fixed size vector, we apply a maximum operation along the rows of 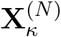. This is typically referred to as a Global Max Pooling layer in neural networks and is repeated *R* times for each row in 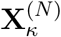:

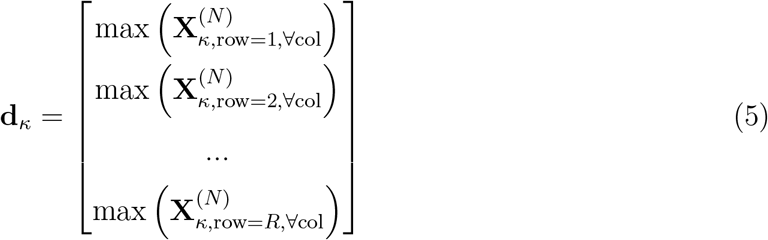

where **d**_*κ*_ is a length *R* column vector regardless of the number of columns in the latent space 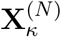. This maximum operation, while important, has the effect of eliminating much of the information in the latent space. To better understand the latent space, we can further process 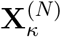 to understand how samples are distributed. For example, a simple operation would be to define another column vector **v** that denotes the total variation in each row of the latent space:

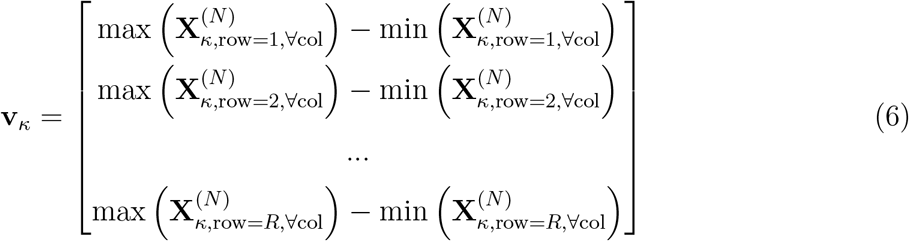

The concatenation of vectors **d** and **v** help elucidate how the samples are distributed in the latent space. As such, we can use these concatenated vectors as inputs to a fully connected dense layer which can learn to interpret the latent space. This output is referred to as the embedding of the protein, **y**_prot_, and is computed as

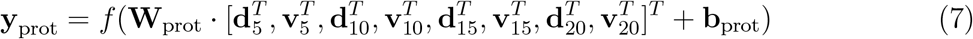

where **W**_prot_ is the learned weight matrix and **b**_prot_ is the bias vector of a fully connected network.

The method described above is similar to a technique recently used in speech verification systems, where the window sizes need to be dynamic because the length of audio snippet is unknown.^51,52^ In speech systems, the latent space is collapsed via mean and standard deviation operations and the embeddings provided for these operations are typically referred to as D-Vectors^51^ or X-Vectors.^52^ In proteins we have a similar problem as the length of the decomposition sequence to consider the active site(s) of protein is dynamic and of unknown sizes. By including the window sizes of 5, 10, 15 and 20 (number of residues to consider at a time), we ensure that the network is able to extract different sized patterns from backbones of varying length.

After embedding the protein and the ligand, we concatenate the vectors together and feed them into the final neural network branch, resulting in a prediction of binding, *ŷ*, which is expected to be closer to “0” for ligands and proteins that do not bind and closer to “1” for proteins and ligands that do bind. This final branch consists of two layers:

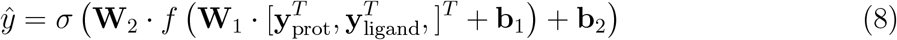

where *σ* refers to a sigmoid function that maps the output to [0, 1]. If we denote the ground truth binding as *y*, which is either 0 or 1, and denote all the parameters inside the network as **W** then the loss function for the SSnet model can be defined as binary cross entropy, which is computed as:

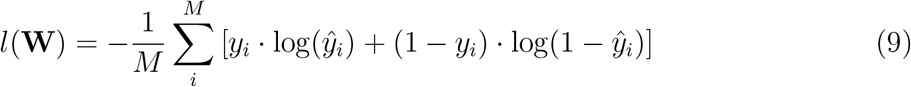

where *M* is the number of samples in the dataset. By optimizing this loss function the neural network can learn to extract meaningful features from the protein and ligand features that relate to binding. At first all weights are initialized randomly and we use back propagation to update the parameters and minimize loss. All operations defined are differentiable, including the collapse of the latent space with Global Max Pooling such that errors in the loss function can back propagate through the network to update all parameters, including the convolutional operations.

The hyperparameters optimized for the model and speed of execution are provided in section 1 of the supporting information.

### Grad-CAM method for heatmap generation

A neural network generally exhibits a large number of weights to be optimized so that complex information can be learned, however, some of this information could be irrelevant to a prediction task. For example, consider the task of identifying if a certain image contains a horse or not. If all horse images also contain a date information on the image and images without horse do not contain date information, the machine will quickly learn to detect the date rather than the goal object (a horse in this case). Therefore, it is essential to verify what a neural network considers “influential” for classification after training. Selvaraju et al. ^36^ proposed a Gradient-weighted Class Activation (Grad-CAM) based method to generate a heatmap which shows important points in the feature data, based on a particular class of prediction. That is, this method uses activations inside the neural network to understand what portions of an image are most influential for a given classification. In the context of protein structures, this methods can help to elucidate which portions of the decomposition plot are most important for a given classification. These influential patterns in the decomposition plot can then be mapped to specific sub-structures in the protein.

Grad-CAM is computed by taking the gradient weight *α*_*k*_ for all channels in a convolutional layer as

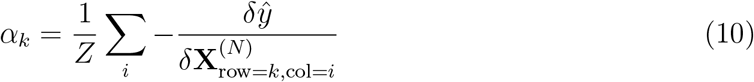

where *k* is the row in the final convolutional layer, *Z* is a normalization term, **X**^(*N*)^ is the activation of the final convolutional layer, and *ŷ* is the final layer output. The heatmap **S** is then computed by the weighted sum of final layer activations:

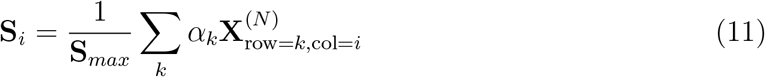

This heatmap **S** specifies the important portions in the input sequence that are most responsible for a particular class activation. For each convolutional branch, we can apply this procedure to understand which portions of the input decomposition sequence are contributing the most, according to each filter size *K* = 5, 10, 15, 20. In this way, we can then map the most influential portions onto locations on the backbone of the protein. To the best of our knowledge, this procedure has never been applied to protein (or ligand) structures because Grad-CAM has been rarely applied outside of image processing.

### Evaluation criteria

The evaluation criteria for PLI are generally presented by the area under the curve of the receiver operating characteristics (AUCROC),^53^ Boltzmann-Enhanced Discrimination (BEDROC), and enrichment factor (EF).^54,55^ AUCROC is primarily used to measure the accuracy of the prediction while both BEDROC and EF measure the early enrichment of true active ligands. To test the accuracy, the receiver operating characteristic curve, which is the plot of true positive rate vs false positive rate, is integrated to get the AUCROC. Thus AUCROC greater than 0.5 suggests that the model performs better than chance. However, AUCROC is not suited for the comparison of models regarding the enrichment of a ranked list with true actives. This problem can be easily illustrated by taking as example two dummy models, A and B. Model A places half of true actives as the top ranking ligands with the other half not recognized as active, while model B randomly ranks the true actives throughout the dataset. In both cases, the AUCROC remains the same, while from a practical perspective, model A is better than model B.^56^

Complementary to AUCROC, EF and BEDROC allow the model to be examined considering its ability to enrich the top ranked ligands. A large number of studies have employed EF to test their models, ^57,58^ and for this reason, values for EF can be easily obtained from the literature. In the present study, only EF is used to compare different models to test the enrichment. EF is defined as

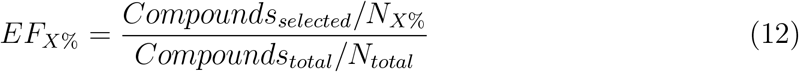

where *N*_*X*%_ is the number of ligands in the top *X*% of the ranked ligands. EF thus considers an estimate on a random distribution for how many more actives can be found withing the early recognition threshold.

### Datasets

Most state-of-the-art models for PLI predictions use human and C. elegans created by Liu et al.^59^ The positive PLIs for these datasets are considered from DrugBank 4.1^60^ and Matador.^61^ The negative PLIs were considered by using ligands for proteins that are dissimilar to the target in query. The human dataset contains 852 unique proteins with at least one positive or negative PLI instance. 1052 unique compounds that bind to these target proteins (one-to-one, one-to-many, and many-to-many) account for 3369 positive interactions. Similarly, C. elegans dataset contains 1434 and 2504 unique proteins and compounds, respectively, for a total of 4000 positive interactions. Experimental setting suggested by Tabei and Yamanishi ^62^ was used such that the ratio of positive to negative interactions used for the training were 1:1, 1:3, and 1:5. A five fold cross validation was performed for evaluation.

Although humans and C. elegans dataset provides good benchmarking against other ML approach for PLI prediction, it does not contain enough PLI instances for use in real world application. Database of Useful Decoys: Enhanced (DUD-E) dataset provides a large number of PLI instances along with computationally generated decoys as the negative PLI instances. More specifically, DUD-E contains 22,886 positive PLIs and 1.4 million decoy over 102 target proteins. The 102 target proteins in DUD-E were divided into 72 and 30 for training and testing, respectively. Each target proteins in DUD-E contains 224 active ligands for each of which 50 decoys that have similar 1D physico-chemical properties, employed to remove bias against dissimilar 2D topology. These decoys are unlikely to bind, and therefore were considered as negative interactions. The net total interactions considered for training were approximately 16 thousand positive PLI and 1 million decoys (negative PLI). In lieu of balancing data, the negative PLIs were dynamically constructed by randomly selecting from 1 million decoys to match the number of positive in each iteration. This trained model is termed SSnet:DUD-E. A schematic representation of the model is shown in Figure S8.

The decoys generated computationally faces the problem of false negatives and therefore an experimental dataset could be more reliable for SSnet. We considered the BindingDB (BDB) dataset^38^ which is a public, web-accessible database of measured experimental binding affinities and contains around 1.3 million data records. We created a database by considering the following properties for each data entry

1. The target has PDB ID cross-referenced as 3D structure. The first annotated structure is taken as reference PDB file.
2. The ligand has SMILES representation in the entry.
3. Record has IC50 value (a measure of strength of binding) and is either less than x (active) or greater than y (inactive).

The values for x assessed were 10 nM, 25 nM, and 100 nM, while the values for y assessed were x nM or 10,000 nM. The preliminary analysis showed that x = 100 nM and y = 10,000 nM provides the best balance between AUCROC and EF_1_% for PLI prediction and as such, this dataset was termed as SSnet:BDB (Figure S11). The dataset contains 4806 unique proteins, and 539,799 (358,023 active and 198,225 inactive) unique PLIs. The dataset was divided similar to DUD-E dataset: 52 proteins for testing and 4754 proteins for training the model. In order to avoid biases due to over-fitting to specific targets, the 4754 proteins considered for training set have less than 75% sequence similarity to the targets in test sets from both DUD-E and BDB datasets. This allows us to confidently test the SSnet:BDB in DUD-E test set. However, the same leniency cannot be applied for SSnet:DUD-E due to its limited target size.

To access an independent dataset, we utilized maximum unbiased validation (MUV) dataset created by Rogers and Hahn ^33^. The MUV dataset is generated from PubChem bioactivity by considering actives that are maximally separated in chemical space to avoid over-representation of physiochemical features. For each target in the MUV dataset a set of decoys was generated with the aim of avoiding analog bias and artificial enrichment. We trimmed the 9 targets as used by Ragoza et al. ^63^ for valid comparison that contains 30 actives and 15,000 decoys for each target.

## Results

Computational methods to predict PLI are often limited by a lack of accurate 3D structures of the regulatory conformation of a target of interest, or by time consuming calculations of diverse protein and ligand conformations. Thus, there is a need for a predictive PLI platform that is both rapid, and can function independent of the target protein conformation. Towards this aim, we have employed an ML/DL based approach (SSnet) based on the curvature and torsion of a protein backbone to develop a predictive PLI algorithm capable of screening 1-billion+ compounds in a manner of days. Specifically, SSnet requires only 18 minutes for computation of one million PLIs to a target using GPU (NVIDIA P100 based on Pascal architecture) accelerated node with Intel Xeon E5-2695v4 2.1 GHz 18-core Broadwell processors and 30 minutes for ten thousand compounds without GPU acceleration. Herein, we first compare SSnet on various computational datasets such as humans, C.elegans and DUD-E. Then we benchmark SSnet by training on completely experimental dataset BDB and compared against state of the art ML/DL PLI algorithms. Lastly, to demonstrate the utility and accuracy of SSnet, we employ the Grad-CAM visualization approach to extract structural features most important to ligand recognition and binding. The Grad-CAM approach both validates the SSnet approach, but can also function as a guide to couple ML-based PLI prediction to downstream analysis using traditional docking-based approaches. Grad-CAM analysis reveals that SSnet can accurately identify regulatory binding sites within protein targets of interest. Importantly, the ability of SSnet to identify these sites is independent of protein conformation, and is able to identify cryptic and allosteric sites without prior information of their regulatory roles. In this manner we demonstrate that SSnet mitigates many of the limitations of alternative predictive PLI approaches, while retaining high accuracy and speed.

### Selection of Ligand Representation

A key bottleneck in development of PLI prediction is the selection of the optimal representation of ligand structure. Several methods of reducing ligand representation to ML/DL methods have been developed. Gómez-Bombarelli et al. ^31^ created a model to generate Continuous Latent Space (CLP) from sparse Simplified Molecular-Input Line-Entry System (SMILES) strings (i.e., a string representation of a molecule) based on a variational autoencoder similar to word embedding.^64^ Scarselli et al. ^34^ proposed a Graph Neural Network (GNN) to describe molecules. Rogers and Hahn ^33^ proposed Extended-Connectivity Fingerprints (ECF), which include the presence of substructures (and therefore also includes stereochemical information) to represent molecules. Riniker and Landrum ^32^ proposed a fingerprint based on substructure and their similarity (Avalon). Since a vectorized representation of protein structure has not been implemented prior to this study, we tested various ligand representations of ligands to find the most suitable and accurate for SSnet. Specifically, we evaluated two traditional ligand fingerprint methods: ECF^33^ and Avalon^32^, as well as two state-of-the-art ML based descriptors: GNN^34^ and CLP.^31^

Table 1 shows the performance of SSnet for the human dataset (1:1 positive to negative) and DUD-E dataset (unbalanced dataset). The GNN descriptor is based on convolution neural networks which require ample amount of data to make sense of the spatial information provided to the model. The descriptor method might also suffer if essential information such as functional groups are deeply embedded in the input data and are not directly accessible to the network. This might be one of the reasons for a lower performance of GNN in terms of AUCROC when compared to ECF and Avalon. CLP gives an AUCROC score of 0.966 and 0.905 for humans and DUD-E datasets respectively. CLP is based on autoencoder which is trained to take an input SMILES string, converts it to a lower dimension, and reproduces the SMILES string back. In this way CLP is able to generate a lower dimensional vector for a given SMILES string. However, relevant information required for the prediction of PLI might not be preserved which explains its low accuracy when used for SSnet. ECF and Avalon have similar AUCROC scores as they both directly provide the information of the atoms and functional groups by considering substructures of a ligand. This implies that the backbone pattern can be best matched with fingerprints that provide substructure and functional group information. We observe the best performance when using ECF, particularly when considering unbalanced dataset of DUD-E.

**Table 1:** Model comparison on the human and DUD-E datasets for various ligand descriptors

| Ligand descriptor | AUCROC |  |
| --- | --- | --- |
|  | human | DUD-E |
| Avalon | 0.982 | 0.968 |
| ECF | **0.982** | **0.974** |
| CLP | 0.966 | 0.905 |
| GNN | 0.944 | 0.972 |

Convolutional neural networks (CNN) have to update a large number of weights and therefore require a large amount of data instances (number of unique PLIs). However, in the human and C. elegans datasets, the number of instances are insufficient, causing SSnet to overfit (Figure S3). To overcome this problem, we ignored the convolution layer and directly fed the proteins’ curvature and torsion to the fully connected dense layer making it similar to ligand vector shown in Figure 2. This helps in reducing the number of weights to be optimized and decreases the chance of overfitting. These approaches were unnecessary for DUD-E since, it contains sufficient instances of data for ML to learn. We note that the approach of removing CNN would still provide a fair comparison of the protein representation compared to other methods. The AUCROC scores are the highest for both humans and DUD-E dataset of 0.982 and 0.974 respectively with ECF, and thus is selected as the ligand descriptor for SSnet.

### SSnet compared on computational datasets

Evaluation of the accuracy of PLI prediction platforms can be complicated by numerous factors, including but not limited to: training set bias as mentioned in Datasets section; lack of true negative instances (computationally generated decoys) as is the case for DUD-E, humans and C. elegans datasets; and, poor comparison metric for end-user, specifically AUCROC which measures overall accuracy without any information pertaining to usability. To provide a clear demonstration of usability, both AUCROC and EF, have been employed to evaluate algorithm accuracy as well as usability. To alleviate any potential bias in SSnet optimization, we trained SSnet on the same dataset as found in the existing literature for direct comparison against models compared. We also retrained the state-of-the-art existing model: GNN-CNN on a larger dataset DUD-E to obtain direct comparisons.

We compared SSnet with PLI specific methods: BLM,^65^ RLS-avg and RLS-Kron,^66^ KBMF2K-classifier, KBMF2K-regression,^67^ and GNN-CNN^27^ with the same experimental setting as Liu et al. ^59^ as shown in Figure 3. It is important to note that BLM, RLS-avg, RLS-Kron, KBMF2K-classifier, and KBMF2K-regression are modeled on properties such as the chemical structure similarity matrix, protein sequence similarity matrix, and PLI matrix. Despite such pre-organized inputs, SSnet was able to outperform in terms of AUCROC (Figure 3). On the other hand, the GNN-CNN model uses a graph neural network for ligands and convolutional neural network for protein sequences. The applicability range of GNN-CNN is superior as it requires only sequence information for a protein compared to SSnet which requires 3D information. However, since the input description of GNN-CNN model limits the model’s capability to extract crucial information embedded in the secondary structure, SSnet outperforms GNN-CNN as demonstrated in Figure 3.

**Figure 3:**
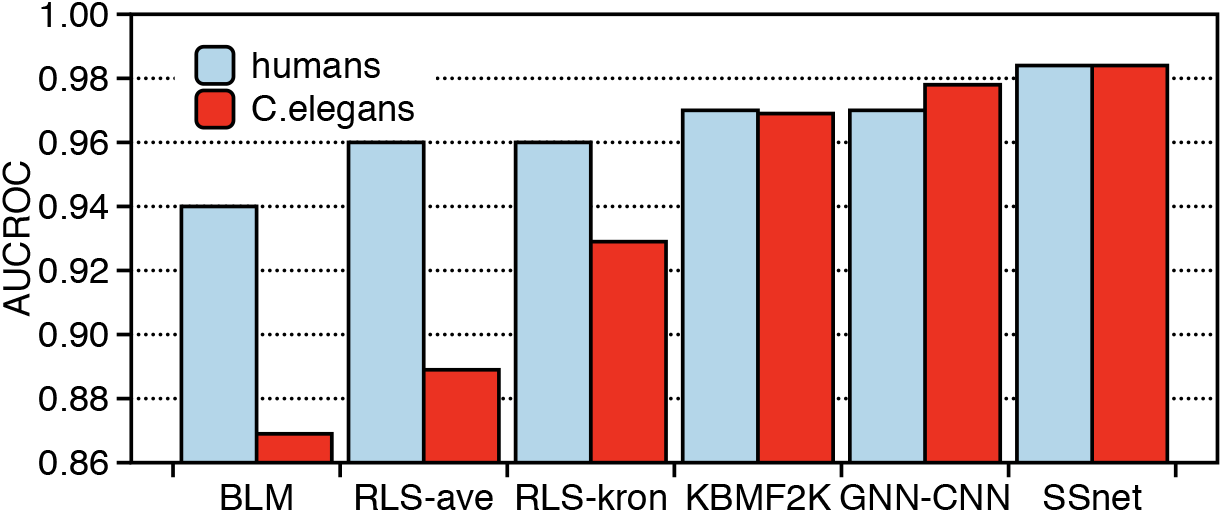
Model comparison on various PLI specific methods with AUCROC. The red color represents SSnet trained on humans dataset and cyan color for C.elegans respectively.

Table 2 shows the comparison of various traditional machine learning models on the human and C. elegans datasets. SSnet outperforms other models in both balanced (1:1) and unbalanced (1:3 as well as 1:5) datasets. This suggests that SSnet is robust and is able to generalize information about the protein and ligand pairs.

**Table 2:** Data comparison (AUCROC) on balanced and unbalanced datasets

| Dataset | k-NN | RF | L2 | SVM | GNN-CNN | SSnet |
| --- | --- | --- | --- | --- | --- | --- |
| humans (1:1) | 0.860 | 0.940 | 0.911 | 0.910 | 0.970 | **0.984** |
| humans (1:3) | 0.904 | 0.954 | 0.920 | 0.942 | 0.950 | **0.978** |
| humans (1:5) | 0.913 | 0.967 | 0.920 | 0.951 | 0.970 | **0.976** |
| C. elegans (1:1) | 0.858 | 0.902 | 0.892 | 0.894 | 0.978 | **0.984** |
| C. elegans (1:3) | 0.892 | 0.926 | 0.896 | 0.901 | 0.971 | **0.983** |
| C. elegans (1:5) | 0.897 | 0.928 | 0.906 | 0.907 | 0.971 | **0.983** |

Note: k-nearest neighbour (k-NN), random forest (RF), L2-logistic (L2) and SVM results were obtained by Liu et al.<sup>59</sup>.

SSnet was trained with the DUD-E dataset, referred as SSnet:DUD-E. The AUCROC is shown in Figure 4 compared against smina, AtomNet, 3D-CNN, and GNN-CNN. The training dataset contains around 16,000 actives and 1 Million computationally generated decoys. Since, we cannot test SSnet against non-ML approaches fairly when trained on a small and limited dataset of humans and C. elegans, DUD-E provides a much fairer dataset for SSnet to compete with the traditional approaches. However, DUD-E dataset is not balanced, and to tackle this issue we trained the model by dynamically constructing balanced datasets. This was achieved by selecting all the actives and randomly selecting equal number of decoys for each iteration. A schematic representation of the model is shown in Figure S8. This procedure helps mitigate any bias that SSnet might have towards a subset of inactives. We compared SSnet:DUD-E with vina^68^ and smina^69^ as traditional docking methods and Atomnet,^26^ 3D-CNN,^63^ and GNN-CNN^27^ as some of the highest performing machine learning models. Figure 4 shows that SSnet:DUD-E outperforms in the average AUCROC score when trained on DUD-E dataset against the most common VS methods available. Similarly, we also compared ML approaches reported for DUD-E dataset. SSnet:DUD-E outperforms Atomnet, 3D-CNN, and GNN-CNN despite using 1D representation of protein structure. Atomnet is a machine learning model that considers vectorized versions of 1 Å 3D grids as input vectors for a protein-ligand complex (Note: Atomnet requires 3D information of protein-ligand complexes). A DNN framework is built based on 3D convolutional layers to predict binary PLI. Similar to Atomnet, 3D-CNN also takes fixed size 3D grid (24 Å) from the centre of the binding site (requires protein-ligand complex) as input which is converted to density distribution around the centre of each atom. These information are then fed to a convolutional neural network to predict PLI. Atomnet and 3D-CNN are based on all atoms in the protein ligand complex. Although a satisfactory information is provided to the model, the large number of input features create noise which makes binary prediction of PLI challenging. With limited amount of information, SSnet was able to outperform all these models in terms of AUCROC with an average score of 0.974. These results suggest that curvature and torsion information accumulates compact information for PLI prediction tasks. The learning curve of loss over epochs is shown as Figure S7a.

**Figure 4:**
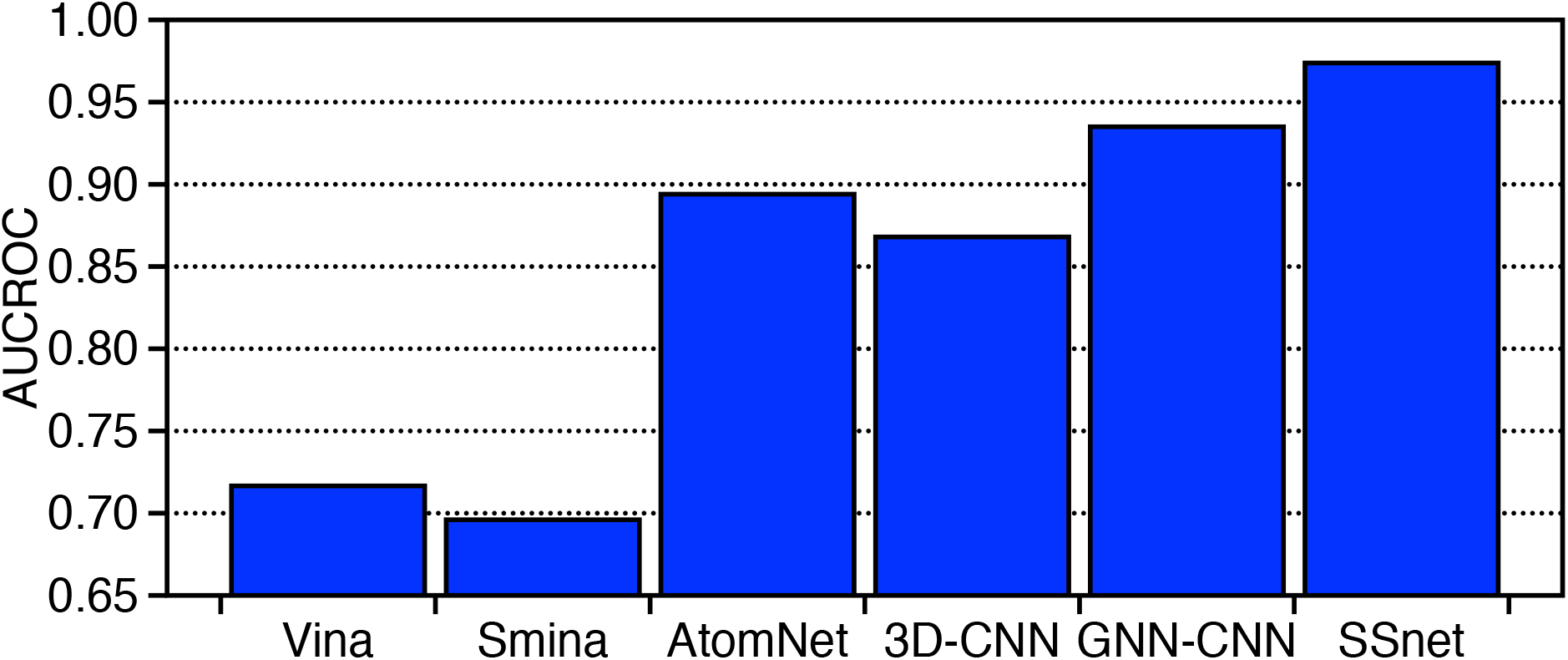
Model comparison of various non-machine and machine learning methods for DUD-E dataset. The AUCROC score for the methods mentioned are derived from the literature.^26,27,63^ SSnet here is trained on DUD-E dataset.

As GNN-CNN is currently the best performing ML/DL for PLI prediction, we compared AUCROC of SSnet by training GNN-CNN (GNN-CNN:DUD-E) following the protocol outlined by Tsubaki et al. ^27^ on the same dataset as SSnet. Vina and 3D-CNN results were obtained by Ragoza et al. ^63^, the result for the four methods are tabulated in Table S6. On DUD-E test set, SSnet:DUD-E performs the best with average AUCROC of 0.97, closely followed by GNN-CNN with 0.96 (Table S6). However, there has been criticism against ML models trained on DUD-E dataset regarding overfitting to the dataset. One of the key criticism of the DUD-E test set is that models trained on DUD-E can easily distinguish active and inactive ligands based on physiochemical properties.^70^ For example, Sieg et al. ^71^ reported that the distributions of MW beyond 500 Da between actives and decoys in DUD-E were mismatched. Further studies have shown that the actives and decoys against the same target can be easily differentiated based on fingerprint.^71–74^ To avoid falling into the pitfalls outlined above, we have tackled these issues by validating the DUD-E trained model using an external dataset. The aim is to show that the features learned by SSnet are not a direct outcome of differences in ligand fingerprints between active and decoys. SSnet:DUD-E is better than GNN-CNN:DUD-E with an average AUCROC of 0.67 versus 0.60 of GNN-CNN:DUD-E (Table S8).

Rogers and Hahn ^33^ generated maximum unbiased validation (MUV) dataset from PubChem bioactivity by considering actives that are maximally separated in chemical space to avoid over-representation of physiochemical features. For each target in the MUV dataset, a set of decoys was generated with the aim of avoiding analog bias and artificial enrichment, two primary causes of overly optimistic predictions in virtual screening. The overall performance across all the targets is essentially random for all methods. The poor performance of various methods over MUV can be attributed to the way MUV creates actives and decoys. Figure S9 shows that SSnet:DUD-E performs equivalent or slightly better than random chance, which is equivalent to or better than the other methods shown. Thus, further discussion of SSnet:DUD-E performance on MUV does not provide any valuable insight.

Table S7 shows the performance of SSnet:DUD-E compared to vina, 3D-CNN and GNN-CNN using the DUD-E test set. The average EF_1%_ over the 21 DUD-E targets were 34, 39, and 41 for 3D-CNN, SSnet:DUD-E, and GNN-CNN:DUD-E respectively for the methods trained on DUD-E dataset. The average EF_1%_ for smina is 8. As an external dataset validation for DUD-E trained model, BDB test set was used shown in Table S8. MUV represents a gold standard for independent validation of PLI prediction. However, for the reasons outlined above, MUV is be an extremely challenging dataset. Similarly to the MUV data described in the section above, we observe all methods perform poorly on MUV (Table S3-5).

### Benchmarking SSnet

BDB is a database of PLIs with reported experimental values for PLI in terms of IC50, EC50, k_*i*_, or k_*d*_. As BDB has large number of PLI instances with annotated experimental values, SSnet can be trained on this dataset while still being able to maximize its learning potential. BDB contains approximately 5000 protein targets, exposing SSnet to a larger portion of the proteomic space and allowing full utilization of convolution network. We benchmark against GNN-CNN, which has outperformed not only partial information (sequence etc.) based model but 3D descriptor models as well, to predict PLI. We retrained and tested GNN-CNN (GNN-CNN:BDB) with the same training/testing used for SSnet (SSnet:BDB). The hyperparameters for GNN-CNN are provided in section 2 of the supporting information. The comparison will give a direct example of whether using secondary structure features improves PLIs prediction over sequence based features.

Figure 5a shows the comparison of ROC curves for prediction of all PLIs in the test set of BDB dataset. SSnet:BDB outperforms GNN-CNN:BDB in terms of AUCROC. Moreover, the ROC curve for SSnet dominates the curve for GNN-CNN:BDB, which supports a conclusion that SSnet is more reliable across all detection thresholds. To test our hypothesis, we performed McNemar test on resultant outputs on the BDB test set from SSnet and GNN-CNN. The MCNemar test statistic observed was 2729.878 and the corresponding p value as 0.0. The test signifies that SSnet outputs are significantly different than that of GNN-CNN. Figure 5b shows the average AUCROC on the test set of BDB. The learning curve of loss over epochs is shown as Figure S7b. SSnet:BDB when tested on BDB test set has the highest average AUCROC of 0.91 which is, followed by 0.85 for GNN-CNN:BDB (Table S8). The scores were based on test set within BDB and thus an independent validation is required to comment on generalizability of the model. We used the DUD-E dataset for this purpose. We eliminated any potential bias from the protein similarity of the test set from DUD-E in the SSnet:BDB training dataset by removing all targets with sequence similarity greater than 75 %.

**Figure 5:**
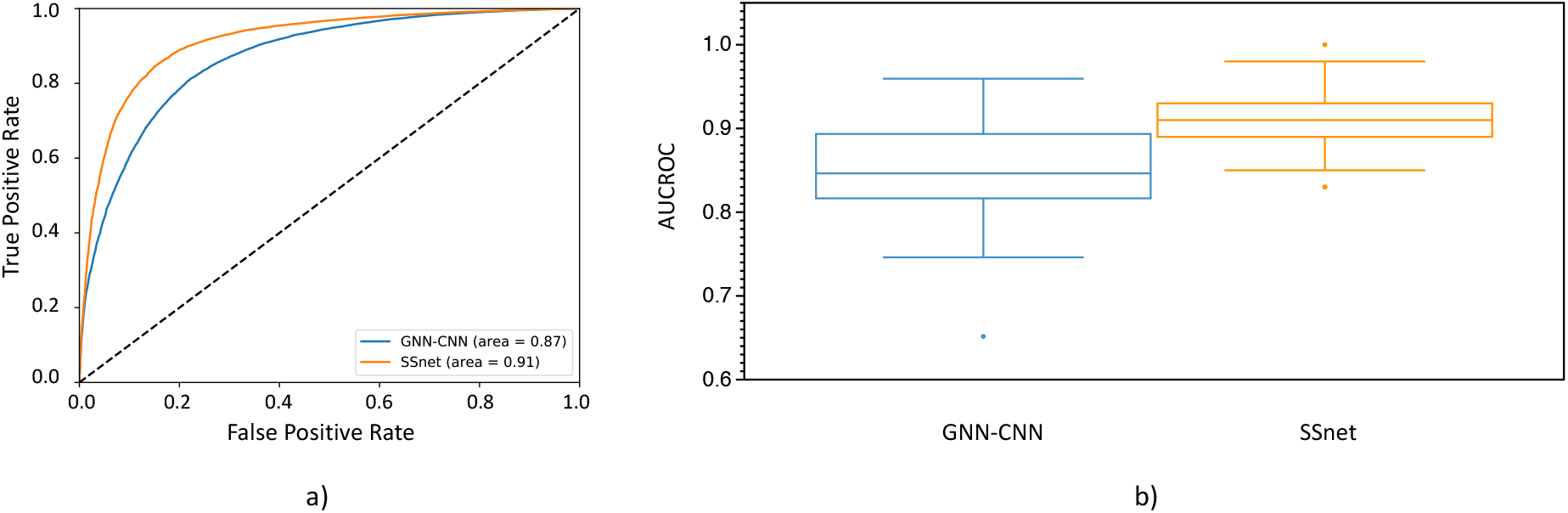
a)ROC plot for the prediction of PLIs using micro-averaging, b) Box and whisker plot for AUCROC for each individual target in the test set of of BindingDB

SSnet:BDB tested on DUD-E has an average AUCROC of 0.81 while GNN-CNN:BDB has 0.79 (Table S6). SSnet:BDB still performs poorly on MUV dataset equivalent to random chance (Figure S9). As SSnet:BDB is based on experimental data, we compared our models with the traditional methods on DUD-E targets. Figure 6a shows the AUCROC of SSnet:BDB compared to the best AUCROC from smina, vina and edock, three freely available traditional virtual screening and docking methods. SSnet consistently has high AUCROC compared to the best scores of smina, vina, and edock.^57^ Figure 6b shows that SSnet:BDB, when tested on the DUD-E dataset, has better performance than random chance (AUCROC greater than 0.5) 94 % of the time with a mean AUCROC 0.78 ± 0.15.

**Figure 6:**
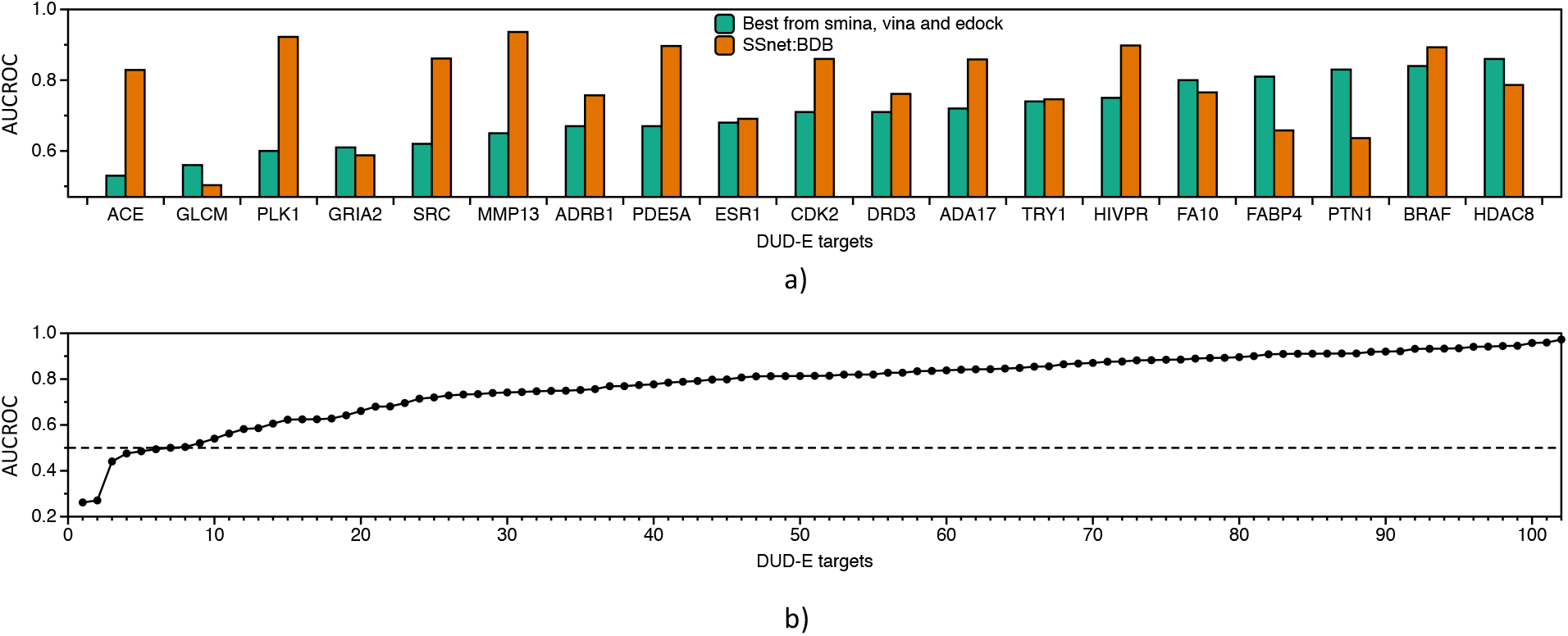
Independent test set DUD-E dataset for BDB model. All targets with more than 75 % similarity in DUD-E dataset were removed from BDB model. a) AUCROC comparison on DUD-E targets over best performer from vina, smina or edock; b) AUCROC comparison with vina for all DUD-E targets.

As described in the evaluation criteria section, AUCROC is not necessarily the best metric for PLI prediction evaluation. While high AUCROC demonstrates the ability of a model in discerning true positives from false positives, this is only useful when selecting drugs with scores greater than 0.5 for SSnet. This might not be feasible when selecting drugs from extremely huge datasets like ZINC which has *1.5 billion ligands. For screening such large databases, enrichment factor (EF) should be employed. In fact, as EF shows the likelihood of finding true actives from the top scoring subset of the database, it provides a more reliable metric for picking high scoring ligands. Furthermore, many studies have shown that ranking based on either metric alone is not sufficient indication of performance.^75–77^

Figure 7a show the EF_1_% of SSnet:BDB compared with the best performance achieved via vina, smina or edock for each target derived from.^57^ SSnet:BDB outperforms the best score of the traditional VS approaches in 74 % of the targets considered. Figure 7b show that for 90 % of the targets, SSnet achieved better outcome than random sampling of the ligands (EF score higher than 1). The average EF_1_% was 15 ± 11.

**Figure 7:**
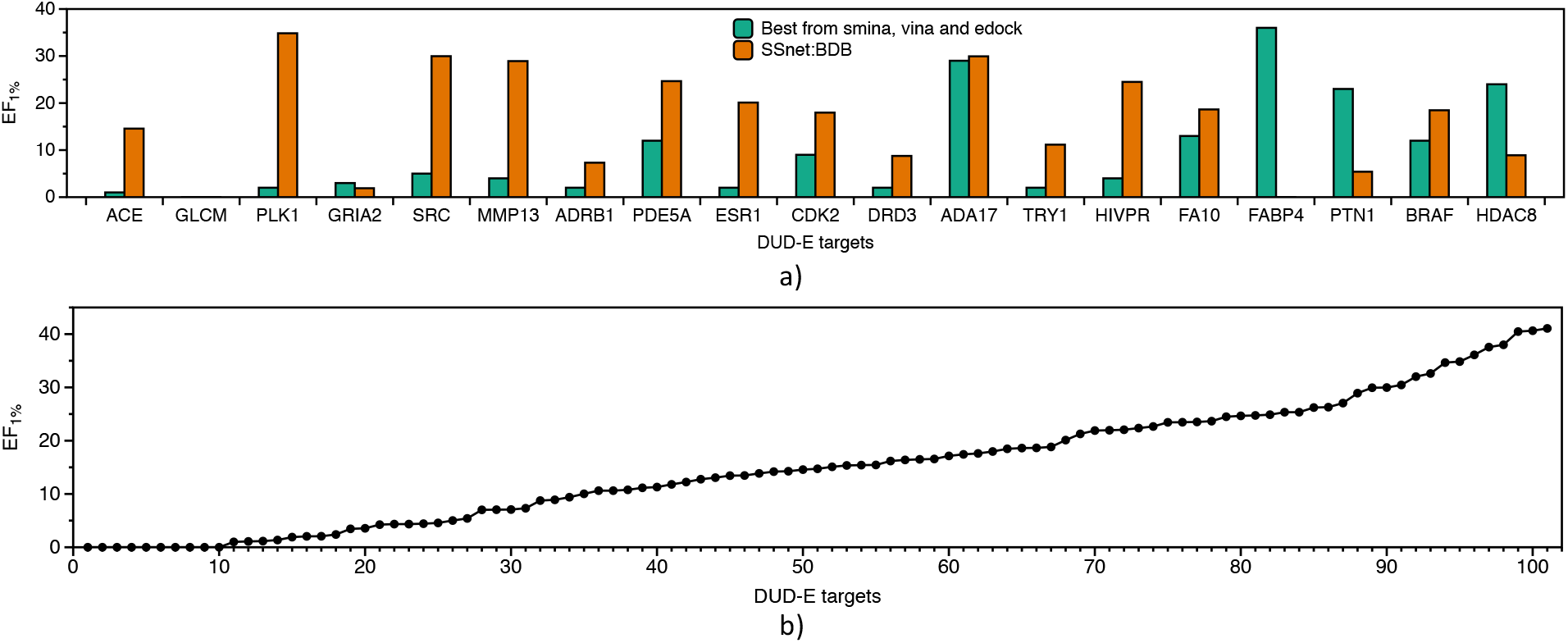
Independent test set DUD-E dataset for BDB model. All targets with more than 75 % similarity in DUD-E dataset were removed from BDB model. a) EF_1%_ comparison on DUD-E targets over best performer from vina, smina or edock; b) EF_1%_ comparison with vina for all DUD-E targets.

SSnet, being an ML/DL model, is not immune to the pitfalls of overfitting. This includes not only removing replicated examples, but also reducing observations with significant similarities. Prior to comparison with advance methods, we made sure to remove possible overlap between BDB and DUD-E dataset using the following protocol:

- Check for ligand similarity by comparing Tanimoto Coefficient (TC) score^78^ for each ligands in BDB train dataset to all ligands of DUD-E dataset.
- Check for fold similarity by comparing TM Score^79,80^ for proteins in the DUD-E dataset to all protein of BDB train dataset

The TC score is a measure of molecular similarity^78^ that compares a distance between the molecular fingerprints and provides a score in the range (0-1] (1 being exactly same). Since we used ECF as the ligand representation, the TC score was determined by considering ECF as fingerprint. Table S9 shows ligand similarity of BDB train dataset to all ligands of DUD-E dataset. We observed that 99.98 % of BDB ligands have maximum TC score of less than 0.85 (for each ligand in BDB train dataset TC scores were computed for all ligands of DUD-E dataset and the maximum TC score was compared). The results signify that there is almost no overlap between the ligands of the two datasets.

To compare the fold similarity between the two datasets, we used TM score.^79,80^ The TM score wights smaller distance errors stronger than larger distance errors, resulting in a normalized score which is length-independent for random structure pairs and sensitive to fold similarity. TM scores are in the range (0-1] and signifies similar folds for > 0.5. Table S10 shows the maximum TM score obtained for each proteins in the DUD-E dataset from all proteins in the BDB train dataset. We observed that 52 % of the DUD-E proteins have similar folds.

Ericksen et al. ^58^ compiled for a set of 21 DUD-E proteins the comparison of industry standard virtual screening methods. We observed that 11 of the DUD-E proteins have similar folds and thus were removed from comparison. The methods used were AutoDock v4.2 (AD4), DOCK v6.7, FRED v3.0.1, HYBRID v3.0.1, PLANTS v1.2, rDock v2013.1, smina 1.1.2 and Surflex-Dock (Surflex) v3.040. Except for AD4 and DOCK which are force field based methods utilizing genetic algorithm and incremental construction respectively, other methods are empirical and knowledge based.^58^ FRED and HYBRID are based on exhaustive rigid docking search. PLANTS uses ant colony optimization, rDock uses genetic algorithm, smina uses iterative local search, and Surflex uses incremental construction by a matching algorithm. Table 3 shows AUCROC for various methods for 21 DUD-E targets.^58^ We observe large variations across different methods for the targets tested on DUD-E dataset. No single method performed the best for all the targets.^81–83^ SSnet:BDB gives an overall superior performance with 4 targets having the best AUCROC score. SSnet:BDB has the highest average AUCROC of 0.81 among the methods shown followed by both FRED and HYBRID 0.78. It is important to note that HYBRID requires and utilizes prior knowledge of the structure of a ligand bound to the target site, which strongly limits the applicability of this method.

**Table 3:** AUCROC comparision on various models

| Target | AD4 | DOCK6 | FRED | HYBRID | PLANTS | rDock | Smina | Surflex | SSnet:BDB | BEST |
| --- | --- | --- | --- | --- | --- | --- | --- | --- | --- | --- |
| ADRB1 | 0.68 | 0.78 | 0.77 | 0.65 | 0.86 | 0.81 | 0.79 | 0.8 | 0.71 | 0.86 |
| DRD3 | 0.69 | 0.59 | 0.79 | 0.81 | 0.69 | 0.66 | 0.68 | 0.71 | 0.73 | 0.81 |
| ESR1 | 0.82 | 0.54 | 0.88 | 0.81 | 0.77 | 0.87 | 0.86 | 0.74 | 0.83 | 0.88 |
| ESR2 | 0.77 | 0.48 | 0.89 | 0.89 | 0.69 | 0.8 | 0.79 | 0.68 | 0.82 | 0.89 |
| ACE | 0.78 | 0.72 | 0.8 | 0.84 | 0.84 | 0.62 | 0.61 | 0.76 | 0.89 | 0.89 |
| HIVINT | 0.54 | 0.65 | 0.74 | 0.6 | 0.76 | 0.67 | 0.81 | 0.66 | 0.50 | 0.81 |
| ADA17 | 0.51 | 0.4 | 0.59 | 0.69 | 0.58 | 0.58 | 0.54 | 0.7 | 0.91 | 0.91 |
| FA10 | 0.86 | 0.81 | 0.79 | 0.82 | 0.8 | 0.9 | 0.84 | 0.76 | 0.90 | 0.90 |
| MMP13 | 0.67 | 0.6 | 0.77 | 0.87 | 0.71 | 0.67 | 0.67 | 0.76 | 0.96 | 0.96 |
| TRY1 | 0.79 | 0.82 | 0.8 | 0.83 | 0.81 | 0.74 | 0.75 | 0.93 | 0.84 | 0.93 |
| mean | 0.71 | 0.64 | 0.78 | 0.78 | 0.75 | 0.73 | 0.73 | 0.75 | 0.81 | 0.88 |
| std. dev. | 0.12 | 0.14 | 0.08 | 0.10 | 0.08 | 0.11 | 0.11 | 0.08 | 0.13 | 0.05 |

Ericksen et al. ^58^ further showed that consensus scoring using the aforementioned methods can boost performance for each target. A consensus scoring is the use of data fusion methods to obtain an improved scoring from the individual scores gathered from various methods. Table 4 shows the scores obtained using various consensus methods applied on AD4, DOCK6 FRED HYBRID, PLANTS, rDOCK, smina and Surflex. The description of the 6 consensus methods are:

**Table 4:**
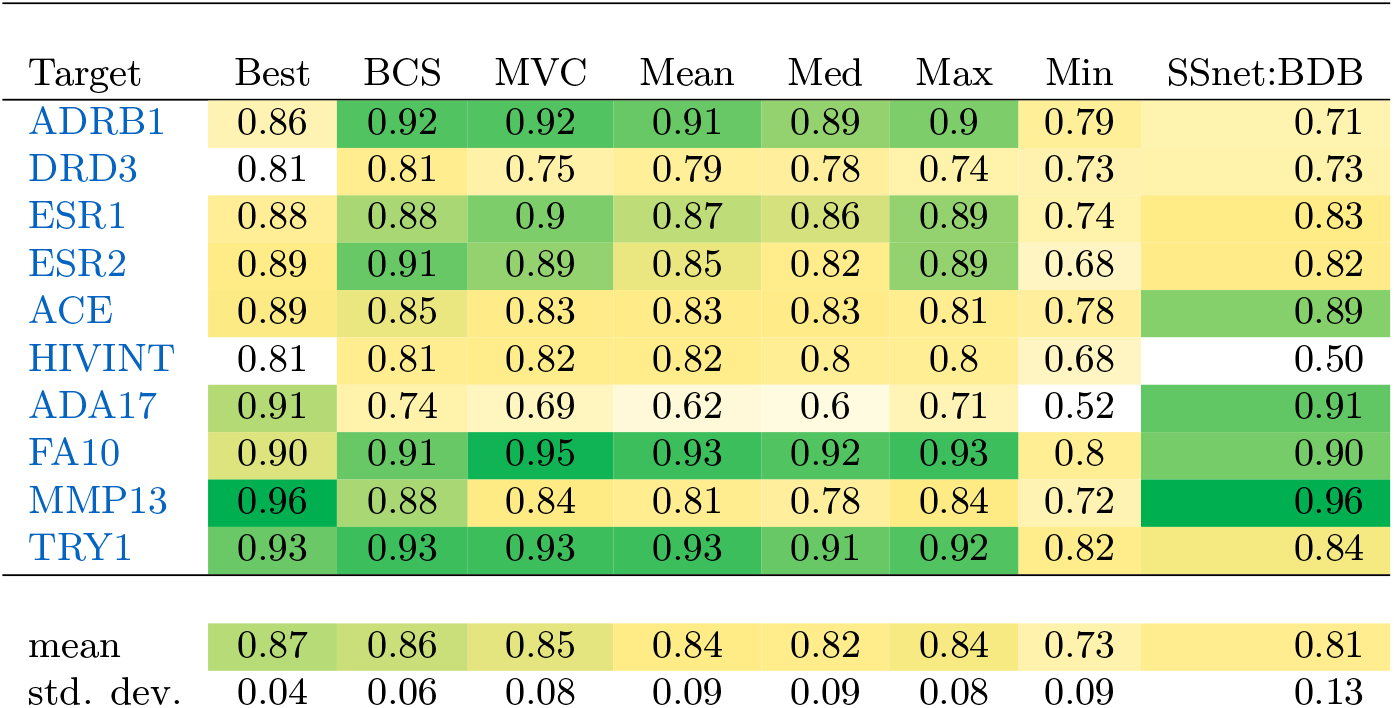
AUCROC comparision on consensus scores

| Target | Best | BCS | MVC | Mean | Med | Max | Min | SSnet:BDB |
| --- | --- | --- | --- | --- | --- | --- | --- | --- |
| ADRB1 | 0.86 | 0.92 | 0.92 | 0.91 | 0.89 | 0.9 | 0.79 | 0.71 |
| DRD3 | 0.81 | 0.81 | 0.75 | 0.79 | 0.78 | 0.74 | 0.73 | 0.73 |
| ESR1 | 0.88 | 0.88 | 0.9 | 0.87 | 0.86 | 0.89 | 0.74 | 0.83 |
| ESR2 | 0.89 | 0.91 | 0.89 | 0.85 | 0.82 | 0.89 | 0.68 | 0.82 |
| ACE | 0.89 | 0.85 | 0.83 | 0.83 | 0.83 | 0.81 | 0.78 | 0.89 |
| HIVINT | 0.81 | 0.81 | 0.82 | 0.82 | 0.8 | 0.8 | 0.68 | 0.50 |
| ADA17 | 0.91 | 0.74 | 0.69 | 0.62 | 0.6 | 0.71 | 0.52 | 0.91 |
| FA10 | 0.90 | 0.91 | 0.95 | 0.93 | 0.92 | 0.93 | 0.8 | 0.90 |
| MMP13 | 0.96 | 0.88 | 0.84 | 0.81 | 0.78 | 0.84 | 0.72 | 0.96 |
| TRY1 | 0.93 | 0.93 | 0.93 | 0.93 | 0.91 | 0.92 | 0.82 | 0.84 |
| mean | 0.87 | 0.86 | 0.85 | 0.84 | 0.82 | 0.84 | 0.73 | 0.81 |
| std. dev. | 0.04 | 0.06 | 0.08 | 0.09 | 0.09 | 0.08 | 0.09 | 0.13 |

**Table 5:**
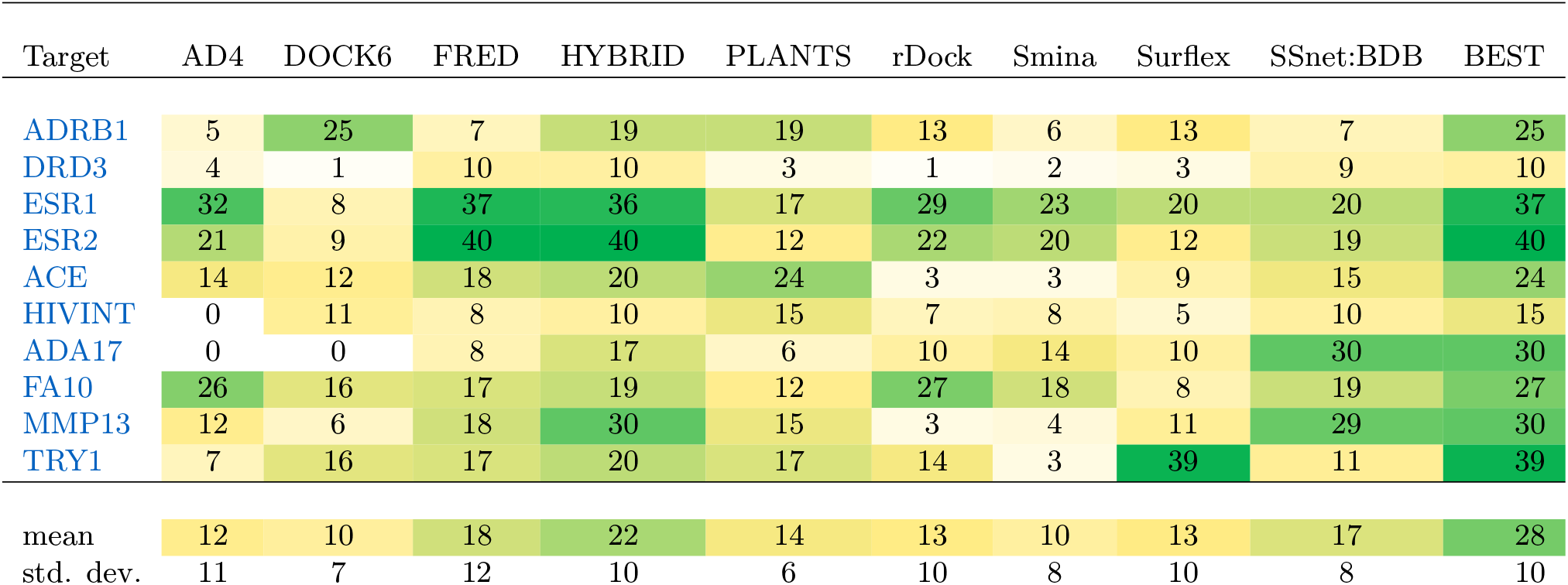
EF_1%_ comparision on various models

- Boosting consensus score (BCS) is a gradient based decision tree framework which is trained on binary labels (actives and decoys) on an individual decision tree for each target where the input is composed of docking scores obtained from each docking method. For each target, the docking scores were provided to other off-targets for boosting the model performance.
- Mean-variance consensus (MVC) is a parameterized function based on gaussian distribution of the scores.
- Mean, median (Med), maximum (Max), and minimum (Min) are the statistics obtained from normalized scores across the docking methods.

The consensus scoring does increase the performance for each target. However, SSnet outperforms or is equivalent to the best consensus based scoring methods for 3 targets in terms of AUCROC. We note that the resources and time required for consensus scoring is significant. SSnet, therefore, serves as a balance between accuracy and resources/time required.

SSnet:BDB has the mean EF_1%_ of 17 signifying that the top 1 % with active ligands on average is 17 times more on average than random picking. SSnet:BDB mean score is similar to most of the methods shown. Only HYBRID and FRED were found to deliver superior performances. Table 6 shows the EF_1%_ obtained using various consensus methods applied on AD4, DOCK6 FRED HYBRID, PLANTS, rDOCK, smina and Surflex. Despite requiring considerably less time and resources compared to consensus methods, SSnet:BDB has high EF_1%_ for most of the targets.

**Table 6:** EF_1%_ comparision on consensus scores

| Target | Best | BCS | MVC | Mean | Med | Max | Min | SSnet:BDB |
| --- | --- | --- | --- | --- | --- | --- | --- | --- |
| ADRB1 | 25 | 31 | 27 | 28 | 24 | 21 | 19 | 7 |
| DRD3 | 10 | 13 | 7 | 12 | 11 | 4 | 11 | 9 |
| ESR1 | 37 | 38 | 37 | 34 | 34 | 32 | 16 | 20 |
| ESR2 | 40 | 35 | 34 | 31 | 28 | 26 | 9 | 19 |
| ACE | 24 | 33 | 30 | 30 | 26 | 19 | 13 | 15 |
| HIVINT | 15 | 19 | 21 | 17 | 11 | 13 | 10 | 10 |
| ADA17 | 30 | 19 | 17 | 16 | 17 | 11 | 4 | 30 |
| FA10 | 27 | 26 | 30 | 33 | 30 | 23 | 19 | 19 |
| MMP13 | 30 | 34 | 25 | 26 | 24 | 20 | 18 | 29 |
| TRY1 | 39 | 33 | 31 | 28 | 27 | 24 | 18 | 11 |
| mean | 30 | 28 | 26 | 26 | 23 | 19 | 14 | 17 |
| std. dev. | 8 | 8 | 9 | 8 | 8 | 8 | 5 | 8 |

### Applicability of SSnet

#### Latent space for proteins

The validation of SSnet demonstrates its reliability. Furthermore, we see that SSnet learns beyond the biases that plague many ML approaches to PLI. Thus, understanding the underlying features learned by SSnet is also of vital importance. To decipher the inner workings of SSnet, we unravelled the global max pooling layer (GMP), shown in Figure 2, using the t-distributed Stochastic Neighbor Embedding (t-SNE). To embed high-dimensional data into low dimension, t-SNE retains similarity information between data points. This allows similar data points in the high dimensional space to form clusters in the lower dimension.

Using SSnet:DUD-E, we tested the proteins in the test set of DUD-E dataset (# of unique proteins = 30) and considered all of their ligand interactions. The results demonstrate that t-SNE clearly distinguishes all proteins (# of clusters = 30) as seen on Figure S7. SSnet:DUD-E had no prior information about the proteins as they were excluded from the training set. The fact that t-SNE clearly distinguishes all the protein suggests that the information gathered by the convolution layers are not general (such as *α* helix or *β* sheet-type patterns) but are specific to PLI. Based on these results, we conclude that our model is able to create a latent space which encodes important information about the bioactivity of the protein. Furthermore, since the model was trained to predict the activity of a protein based on several ligands, such latent space will encode important information about its binding site and therefore can be a powerful tool to compare proteins based on their activity. To identify the protein features that SSnet considers, we performed Grad-CAM analysis described in the section below.

#### Visualization of heatmap using Grad-CAM

ML/DL models have large number of weights that are optimized to learn complex information, and therefore it is important to investigate which input features are critical for the learning process of the model. In most of the previous studies, ^27^ a neural attention layer is added to the network to understand the important pathways in the feature space that get higher attention relative to others. However, the information learned from a neural attention layer could be misinterpreted since it adds an additional layer, increasing the complexity of the network. To tackle this problem we opted for Grad-CAM, since it can provide an insight into the activated pathways without adding complexity to the network. These activated pathways can then be traced back to the input features that are important in predicting a particular class based on convolution outputs.

Grad-CAM highlights the important residues for ligand recognition. In all cases the ligand forms several types of PLIs, some of which were analyzed as shown in Table 7. Analyzing the highlighted structure from the Grad-CAM analysis, we observed that the SSnet considers a weighted probability density of the binding sites present in a protein (Figure S10). It is important to note that the analyzed proteins include allosteric sites, an example of which is shown in Figure 8. The protein Prolyl-tRNA Synthetase from *Plasmodium falciparum* is in complex with glyburide. Hewitt et al. ^84^ showed that glyburide binds to the allosteric site of Prolyl-tRNA Synthetase. SSnet is able to highlight the region of the protein where glyburide binds, which is not the known orthosteric binding site but an allosteric one. This information can be used by researchers to describe bounding box for any downstream docking application.

**Table 7:** Percentage of detected residues by SSnet

| Cutoff<br>( in Å ) | Truly detected<br>residue | Covalently<br>involved | Electrostatically<br>involved | Hydrogen bond<br>involved | Metal Ligand |
| --- | --- | --- | --- | --- | --- |
| 0 | 50.4 | 38.1 | 51.2 | 45.1 | 67.2 |
| 4 | 69.0 | 57.9 | 70.0 | 64.8 | 82.5 |
| 6 | 81.7 | 78.5 | 80.1 | 80.3 | 88.6 |
| 8 | 89.1 | 93.2 | 88.3 | 88.0 | 93.0 |
| # of annotated<br>residues | 11936 | 354 | 2867 | 3436 | 1665 |

**Figure 8:**
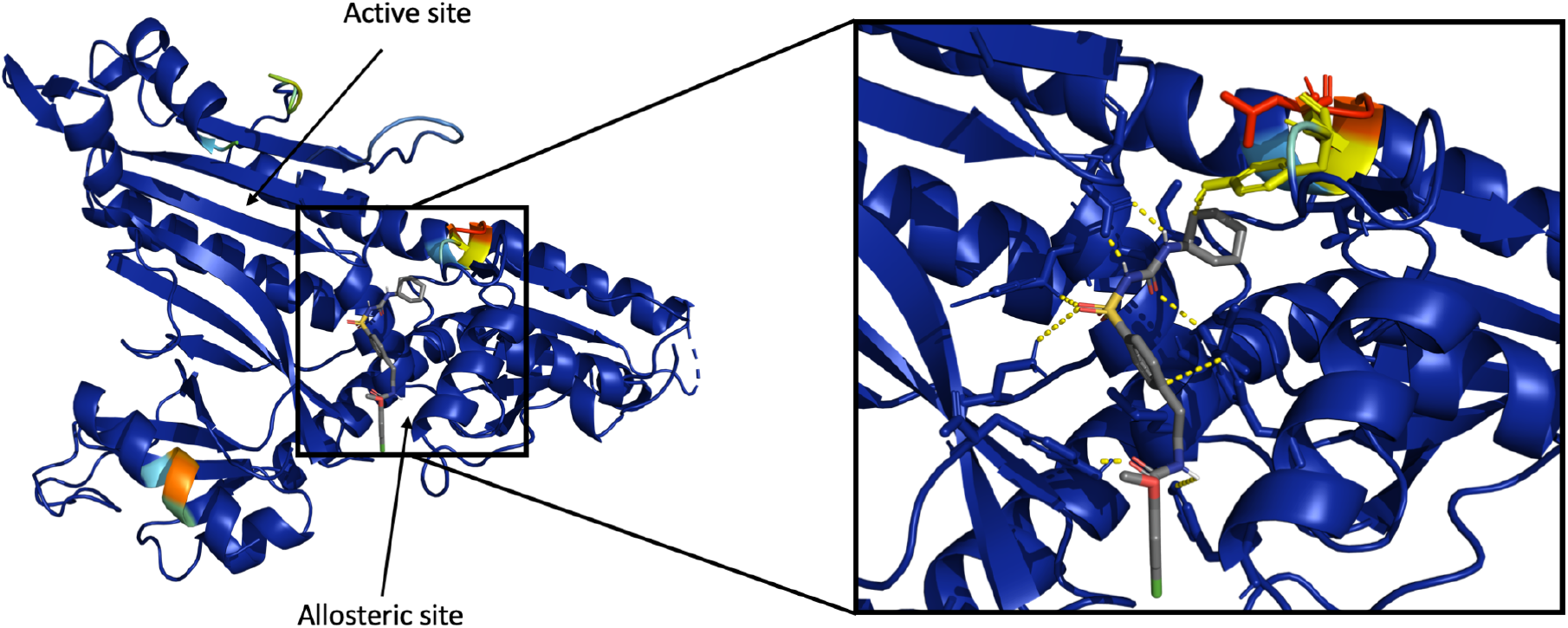
Grad-CAM visualization of heatmap for the protein Prolyl-tRNA Synthetase. The heatmap is a rainbow mapping with violet as the lowest and red as the highest value.

The interaction between proteins and ligands can be mediated through both covalent and non-covalent interactions. Furthermore, various non-covalent interactions that participate in PLIs, including but not limited to hydrogen bonding, Van der Waals interaction, and electrostatic interaction, have been identified. To decipher if SSnet is learning general information critical for PLI, we conducted a test on The Catalytic Site Atlas (CSA) database.^85^ The CSA database contains enzyme active sites and annotated catalytic residues. The entries are either hand-annotated which are derived from literature or homologous entries based on sequence similarity. Proteins with problematic structures, such as extremely large structures (more than 2500 residues per chain) or missing a large chunk of residues (more than 30 missing residues), were removed. This resulted in 577 unique proteins and 11,936 unique annotated residues.

From the Grad-CAM highlighted structures, we identified the residues that maximally influenced PLI prediction for SSnet. These residues were then cross-referenced to CSA. Table 7 shows the percentage of residues detected by SSnet in close proximity at various cutoff lengths for each annotated residue. At 0 Å cutoff distance SSnet correctly detected 50 % of the annotated residues. Specifically, 38, 45, 51 and 67 % of annotated interactions that involve covalent bond, hydrogen bond, electrostatic interaction and metal ligand respectively were identified from the complete set. We observe 89 % of the annotated residues within 8 Å of the highlighted region. It is important to note that these annotated residues envelop various binding sites such as allosteric, cryptic, catalytic etc. The overall trend for individual interaction follows similar to all annotated residue detection. The result shows that SSnet extracts fold information required for PLIs while remaining unbiased towards any particular type of interaction. PLIs predicted are a consequence of multiple factors deeply embedded in the fold of the protein. To further investigate how SSnet processes protein folds information, we tested the applicability of SSnet to identify PLI independent of the structure conformation.

#### SSnet is conformation blind

The binding of a ligand perturbs the secondary structure and can cause significant differences from the original unbound protein structure. We investigated the ability of SSnet in predicting ligand binding based on an unbound protein structure. We divulged our focus on answering two key questions:

- Can SSnet predict the same results using an unbound protein structure or a different conformation of the same protein?
- Can SSnet detect cryptic sites based on unbound protein structures?

To address the first question, Table 8 shows the results of binding site prediction when different protein conformations of 9 randomly selected targets from the test set of DUD-E dataset. Each target was screened through 45,609 randomly selected ligands from DUD-E dataset. The first and second columns denote the PDB ID for a protein ligand complex (PLC) in the DUD-E dataset and PDB ID of a different conformation (DC) of the same protein, respectively. The first five rows have DC with the same protein in PLC bound with a different ligand, and the remaining are apo proteins (unbound proteins) of the PLCs. The presence of a ligand changes the secondary structure of the protein and therefore we observe a range of root-mean-squared-distance (RMSD) from 0.175 to 0.666 between a PLC-DC pair. The prediction results for each PLC-DC pair in predicting actives and inactives are almost the same with maximum error of 0.03 %. To analyse further we looked into the probability scores obtained for each ligand. The Figure S12 shows the correlation of SSnet scores for two conformations of same proteins plotted against each other. The plots demonstrate the conformational blindness of SSnet. Conformational blindness of SSnet can be attributed to the convolution network learning the fold patterns required for PLI. This success highlights the robustness of representing the proteins in terms of torsion and curvature. Torsion and curvature are sufficient in representing subtle changes in the local fold. This further suggests that SSnet is able to predict similar results for a given protein regardless its specific conformation.

**Table 8:** Comparison on performance of SSnet on different conformations of a protein

| PLC<sup>a</sup> | DC<sup>b</sup> | State<sup>c</sup> | RMSD<sup>d</sup> | Actives |  | Inactives |  | Error<sup>e</sup> |
| --- | --- | --- | --- | --- | --- | --- | --- | --- |
|  |  |  |  | PLC | DC | PLC | DC |  |
| 1B9V | 1B9S | Bound | 0.267 | 22518 | 22518 | 23091 | 22518 | 0.00 % |
| 1C8K | 8GPB | Bound | 0.279 | 22505 | 22498 | 23104 | 23111 | 0.03 % |
| 1MV9 | 1MVC | Bound | 0.175 | 22434 | 22436 | 23175 | 23173 | 0.01 % |
| 1Q4X | 2J4A | Bound | 0.463 | 22507 | 22509 | 23102 | 23100 | 0.01 % |
| 1QW6 | 1QWC | Bound | 0.237 | 22507 | 22509 | 23102 | 23100 | 0.01 % |
| 1BCD | 2FNM | Unbound | 0.270 | 22518 | 22518 | 23091 | 23091 | 0.00 % |
| 1H00 | 4EK3 | Unbound | 0.178 | 22518 | 22518 | 23091 | 23091 | 0.00 % |
| 1J4H | 5HT1 | Unbound | 0.666 | 22518 | 22518 | 23091 | 23091 | 0.00 % |
| 1KVO | 1MF4 | Unbound | 0.565 | 22517 | 22521 | 23092 | 23088 | 0.01 % |

<sup>a</sup>Protein-ligand complex from test set of DUD-E dataset
<sup>b</sup>Different conformation of PLC
<sup>c</sup>Bound refers to DC with a different ligand and unbound refers to DC with no ligand
<sup>d</sup>Root-mean-squared distance between PLC and DC
<sup>e</sup>Percentage error in predicting actives
\*All PLC and DC rows refer to 4 letter PDB ID.

Some proteins have binding sites that are not easily detectable. These proteins, termed cryptic proteins, have binding sites that are present in a protein-ligand complex crystal structures but not necessarily in the apo protein crystal structures.^86^ The change in conformation upon ligand binding is a dynamic phenomenon and has been widely reported in the literature.^87^

Figure 9 shows cryptic sites for 3 different proteins taken from CryptoSite set. ^88^ Figure 9 shows the bound (proteins with heatmap) and unbound (grey) proteins. The result from Grad-CAM analysis of these proteins are highlighted with blue having the lowest and red having the highest influence on the PLI prediction. Furthermore, we notice that SSnet score was not as strongly influenced by the residues as is the case for the proteins shown in Figure 8. This is visually observed by higher abundance of intermediate colors (yellow-green) in the cryptic sites (Figure 9, compared to mostly red or blue in Figure 8. Figure 9a shows the unbound-bound pair of a cAMP-dependent protein kinase. The unbound structure of this protein (PDB-ID 2GFC) has an activation loop that protrudes into the active site, occluding the binding pocket. SSnet is able to predict that the ligand will bind strongly to this protein and highlights location closer to the actual binding site on the unbound structure, even though in the latter the binding site is occluded. Retrieving such information is of critical importance as these sites are practically impossible to detect using classical VS methods as they rely on the particular protein structure used for the calculation. Figure 9b shows the bound-unbound pair for Tyrosine kinase domain of hepatocyte growth factor receptor C-MET from *Homo sapiens* (PDB ID 3F82 and 1R1W, respectively).

**Figure 9:**
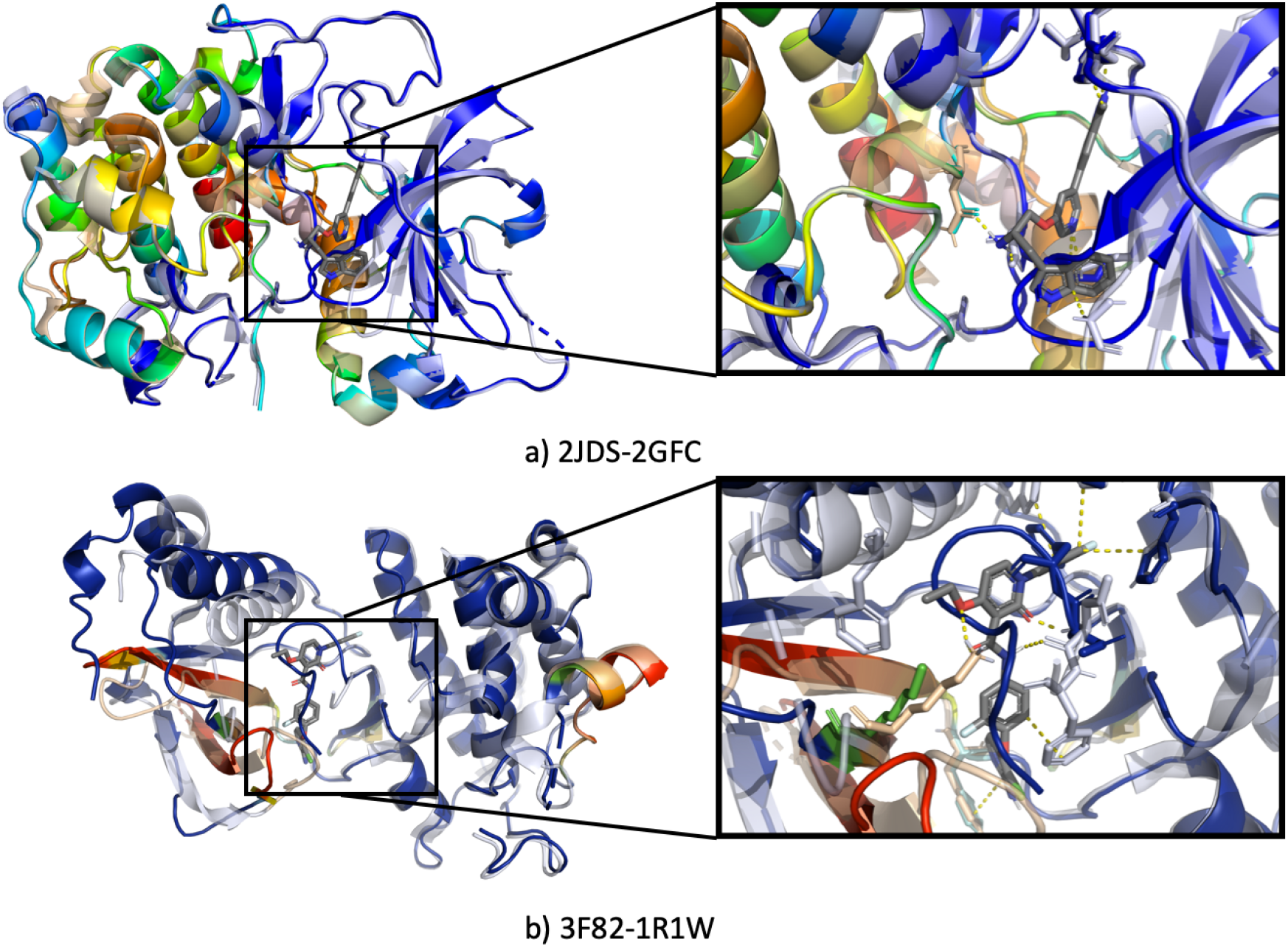
Heatmap generated from unbound protein for cryptic sites. The heatmap is a rainbow mapping with violet as the lowest and red as the highest value. The grey color shows bound protein with ligand.

As the ligand binds at flexible regions in the protein, we observe large conformation changes that result in the formation of a pocket for ligand binding. This example shows that the predicted binding is a consequence of each individual chain, considered independently. Thus, the applicability of SSnet can be expanded for predicting PLIs that involve multiple protein chains.

Grad-CAM analysis, as well as conformation independence of PLI prediction, shows that SSnet learns crucial details of the input features required for predicting PLI. The torsion and curvature of the protein structure effectively describe the features required for PLI while remaining compact, as it is a vectorized format of the complex protein structure. The ability of SSnet to predict PLIs regardless of the crystal conformation showcases the versatile nature of the model. Grad-CAM analysis of the PLI prediction enhances the result of the screening as it can be quickly paired with pre-existing docking tools that require rigid bounding box for accurate posing and subsequent docking.

## Discussion

With our analysis it seems that protein folds play one of the key factor in PLI. This is highlighted by the high accuracy of SSnet and the grad-CAM analysis where we observed that regions near the binding pocket were most influential for the prediction task (89 % accurate withing 8 Å of a residue), signifying fold dependency of a ligand. It is important to note that SSnet had no prior information about the binding pocket. The claim that a molecule should have lower than 500 db as molecular weight to be drug-like,^89^ further shows the dependency of ligands on protein folds. Concerning the complex involvement of ligands in PLIs, a hypothesis that a protein fold holds information about the potential interactions that might be induced though the protein side chains in the binding pocket can be inferred. However, SSnet being blind to conformation limits its capability to account for mutations resulting to the same fold but significant difference in binding affinity. Thus, SSnet should be treated as a first hand screening tool to cull millions of drug-like molecules and not as an exact binding affinity prediction method. Further validation utilizing high accuracy docking methods, molecular dynamics simulations or experimental validation would be of critical importance.

## Conclusion

The study of PLI is an important field for progress in pharmaceutical industry and potentially extended to any biological applications. The limitations of the existing tools for predicting PLI, however, has stalled the progress in these fields. PLI computations suffer from large compute times as accurate PLI prediction require accounting of large number of physico-chemical properties. Furthermore, biases arise for the ML based PLI prediction tools due to imbalances in the representation of these physicochemical properties in the training dataset. On the other hand, classical methods rely on stochastic optimizations such as Monte-Carlo or genetic algorithm type approaches to generate the poses for the ligands and subsequently minimize the bound structures. These approaches require precise 3D conformation and have much higher computing times. SSnet does not show biases in physicochemical properties and necessity of accurate 3D conformation while requiring significantly less computing time. This is achieved by utilization of secondary structure information in the form of curvature and torsion of the protein backbone. The ML model employed enables fast computation once the model is trained as once the weights are fixed, prediction is result of multiple subsequent matrix transformation. The CNN framework employed enables SSnet to learn PLI patterns across wide range of residue interactions, encrypted into the torsion and curvature of the protein backbone. The overall architecture of SSnet therefore works in tandem to eliminate biases that plague many other ML approaches while retaining the speed.

SSnet outperforms several notable ML algorithms in terms of AUCROC when trained and tested on humans and C. elegans dataset. Furthermore, SSnet outperforms the state-of-the-art ML approach GNN-CNN in terms of AUCROC and EF1% when using both models trained on DUD-E and BDB datasets. This comparison holds true even when both the models were tested on independent test sets. Moreover, SSnet:BDB performs better if not equivalent to the classical methods in terms of both AUCROC and EF while being orders of magnitude faster than any of the traditional VS approaches.

The SSnet model utilizes secondary structure information of the protein and, since it only processes CA atoms, it does not necessarily require high resolution structural information. The analysis done on bound-unbound proteins show that SSnet can predict similar results even with different conformations of the protein including cryptic sites. SSnet requires a single conformation to predict whether a ligand is active or not, even if the protein-ligand complex has different tertiary structure. Grad-CAM analysis not only addressed the validity of SSnet learning appropriate details from the input features but also provides the user an intuitive visualization of potential binding site for PLI.

SSnet can be coupled with traditional VS/docking algorithm as pre-screen to filter ligands. Moreover, Grad-CAM analysis showed that SSnet is able to provide accurate prediction of ligand binding sites: active, allosteric, and cryptic sites. As most of VS/docking algorithms necessitate prior knowledge of the binding site, this information can be used to trim ligand search space and determine the box placement. Such information is not retrievable by most of the other VS methods for PLIs prediction. Furthermore, a standalone package has been provided for Linux, Windows and OS-X, making it readily available to users at all levels of computational expertise and not just users familiar with programming. These features of SSnet allows it to be seamlessly integrated into existing VS workflow where SSnet can be used to cull large databases to a small size, and determine bounding box for subsequent docking algorithms.

The top scoring docked poses can then be used directly in experimental setup or further analysis using techniques like Molecular Dynamics, to study the PLI.

Our study suggests that end-to-end learning models based on the secondary structure of proteins have great potential in bioinformatics which is not just confined to protein ligand prediction and can be extended to various biological studies such as protein-protein interaction, protein-DNA interaction, protein-RNA interactions etc. Inspired by the t-SNE results for the last layer in protein embedding we propose a possible latent space for proteins that encodes important information about the protein bioactivity and further exploration could result in a metric to compare proteins based on their bioactivity. We leave these explorations of both the SSnet model and the underlying latent space for future work.

To ensure replicability of both model generation as well as model validation, all scripts developed and implemented in this work are provided through GitHub (https://github.com/ekraka/SSnet) under MIT License without any restrictions or liability.

## Acknowledgements

The authors thank SMU for generous supercomputer resources. The authors would also like to thank **Saeedi Mohammadi** for the discussion of SSnet and providing helpful tips.

## Graphical TOC Entry

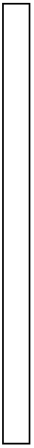

