## Supplemental information for "SSnet: A Deep Learning Approach for Protein-Ligand Interaction Prediction"

#### SSnet model

The SSnet model was tested on SMU Maneframe II. The GPU node is equiped with 36 accelerator nodes with NVIDIA GPUs, dual Intel Xeon E5-2695v4 2.1 GHz 18-core "Broadwell" processors, 256 GB of DDR4-2400 memory, and one NVIDIA P100 GPU accelerator. Each NVIDIA P100 GPU has 3,584 CUDA cores and 16 GB CoWoS HBM2 memory. The P100 GPU is the based on the new Pascal architecture and an extremely high bandwidth (732 GB/s) stacked memory architecture. In the following we discuss the hyper-parameters inlovlved in the optimization of the model.

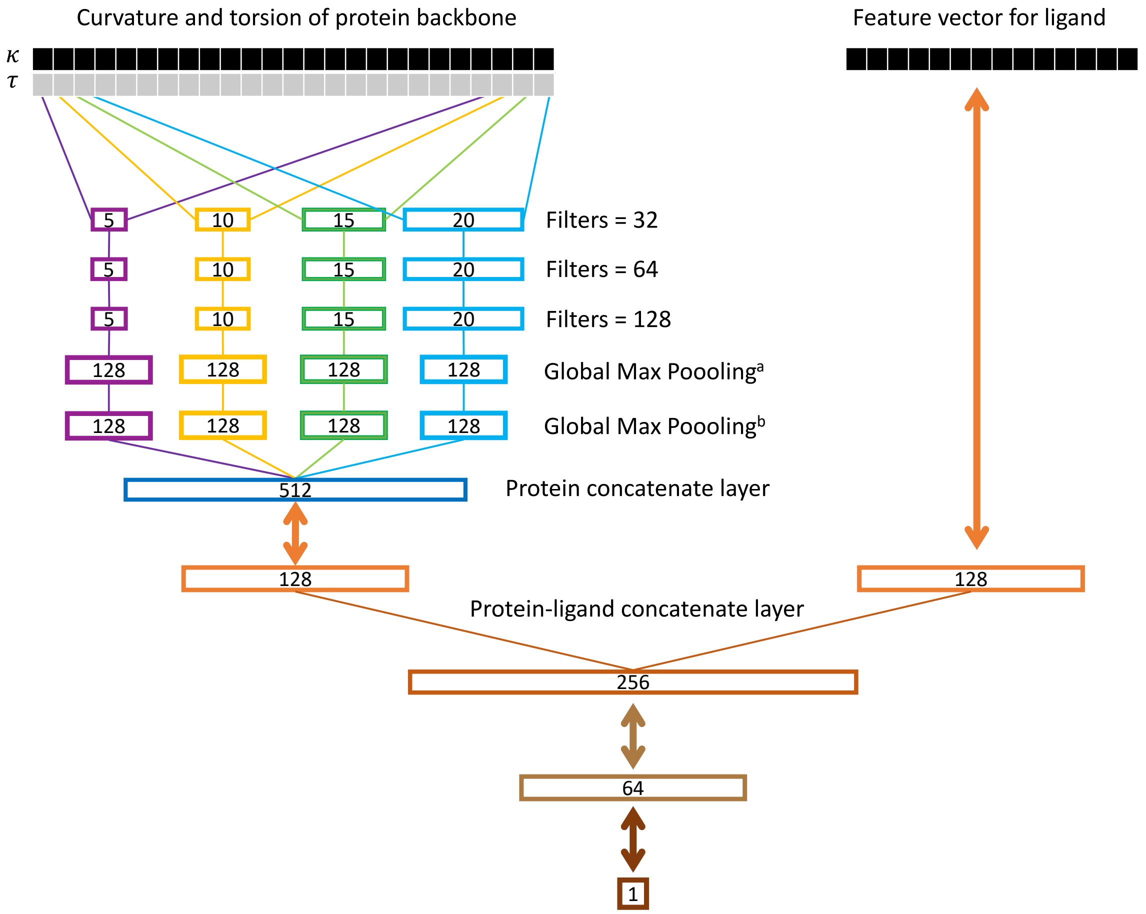


Figure 1: SSnet model. The curvature and torsion pattern of a protein backbone is fed through multiple convolution networks with varying window sizes as branch convolution. Each branch further goes through more convolution with same window size (red, orange, green and light blue boxes). A global max pooling layer is implemented to get the protein vector. The ligand vector is directly fed to the network. Each double array line implies a fully connected dense layer. The number inside a box represents the dimension of the corresponding vector.

#### Layer Width

The increase in layer width significantly reduces time, however, the loss remains almost constant. We came up with the optimized parameters as shown in Figure 1.

#### Model Depth

Model depth behaved similar to layer width. We observed a depth of three layers deep is optimal in performance compared to loss.

#### Pooling Type

We used two global max pooling layers. More details about the pooling layers are discussed in the main text. In sort using two global max pooling layers we tend to capture the variance of the data.

#### Fully Connected Layer

Modifications to the final fully connected layer had no discernible effects on predictive performance or training time, suggesting most of the learning is taking place in the convolutional layers.

#### Time comparisons

SSnet model takes on an average 332 seconds per epoch to train on DUD-E dataset (63,120 instances) and 1178 seconds per epoch on BindingDB dataset (233,573 instances).

#### Data augmentation

Drug augmentation is a technique to increase number of instances to train by adding some sensible noise from the train data. The SSnet model takes up 6 different chains arranged in a stack of 1500 amino acids each (padded with zeros). The chains were randomly shuffled so that the model does not rely on the ordering of chains.

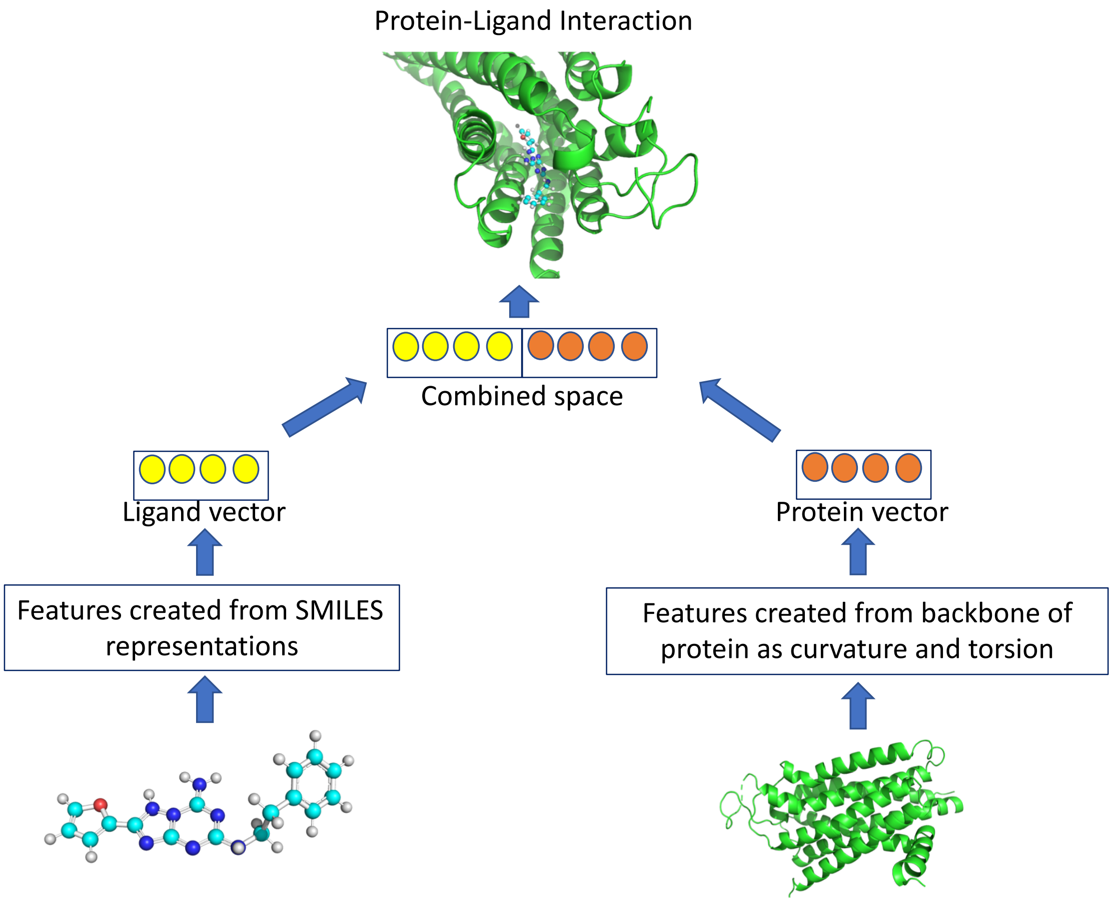


Figure 2: SSnet model overview. The SMILES string and the PDB file for ligand and protein respectively is fed to the model which is converted to ligand vector and protein vector respectively. The two vectors are then concatenated and fed to further networks for PLI predictions

Table 1: Model comparison on the DUD-E dataset for various ligand descriptors

| Ligand descriptors | AUC |
|---|---|
| GNN | 0.983 |
| ECF | 0.984 |
| CLP | 0.906 |
| Avalon | 0.964 |

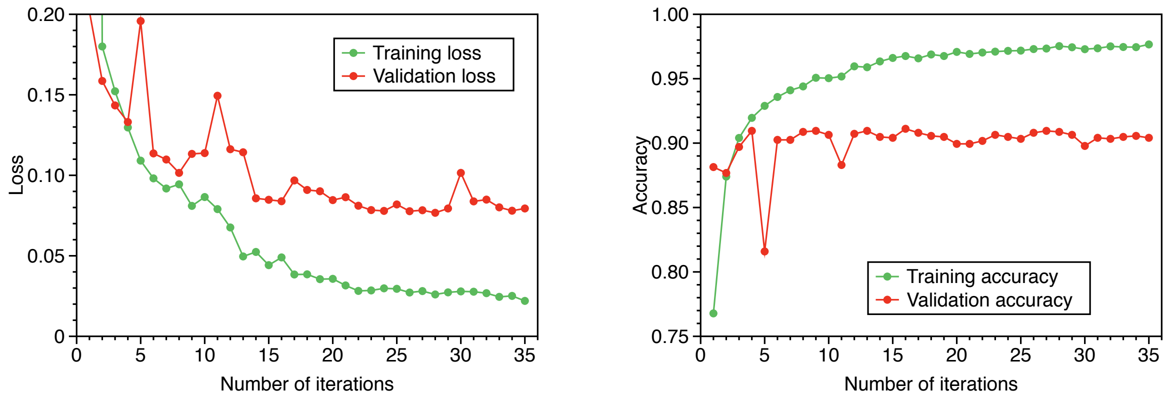


(a) (b)

Figure 3: SSnet model overfits when convolution neural network is applied to smaller datasets such as human or C.*elegans*

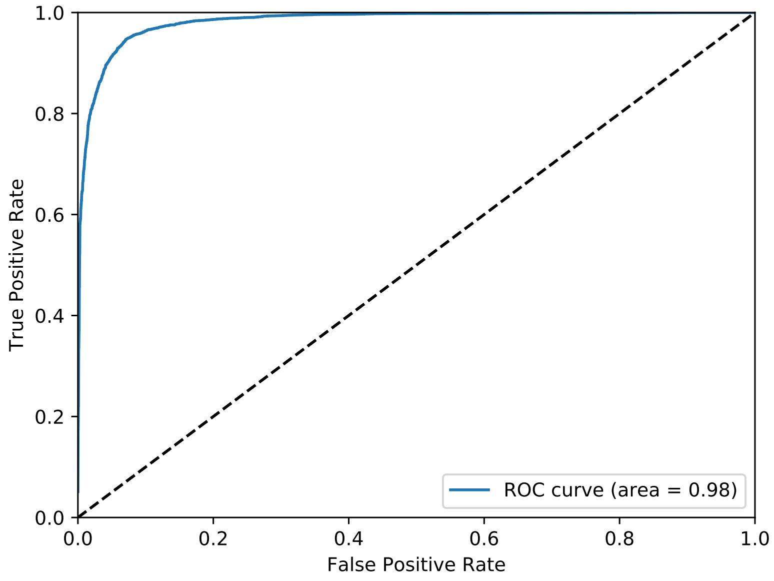


Figure 4: Receiver operating characteristics for the predictions on DUD-E dataset.

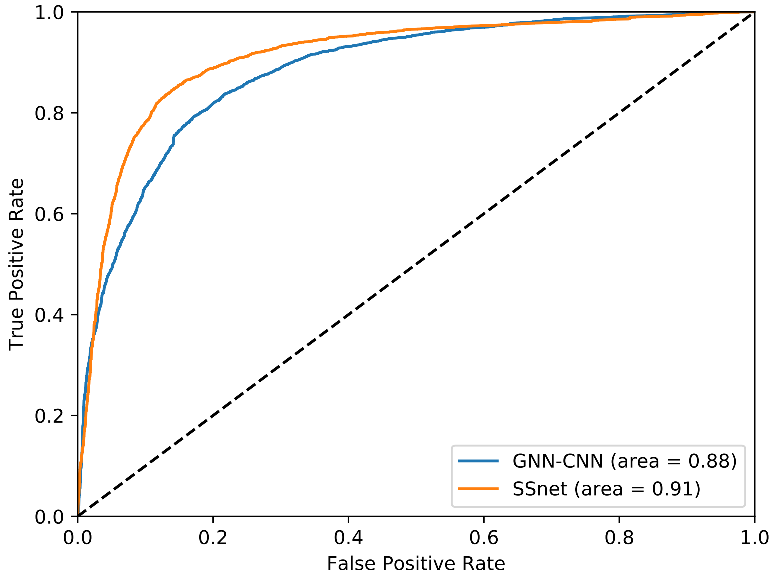


Figure 5: Receiver operating characteristics for the predictions on BindingDB dataset.

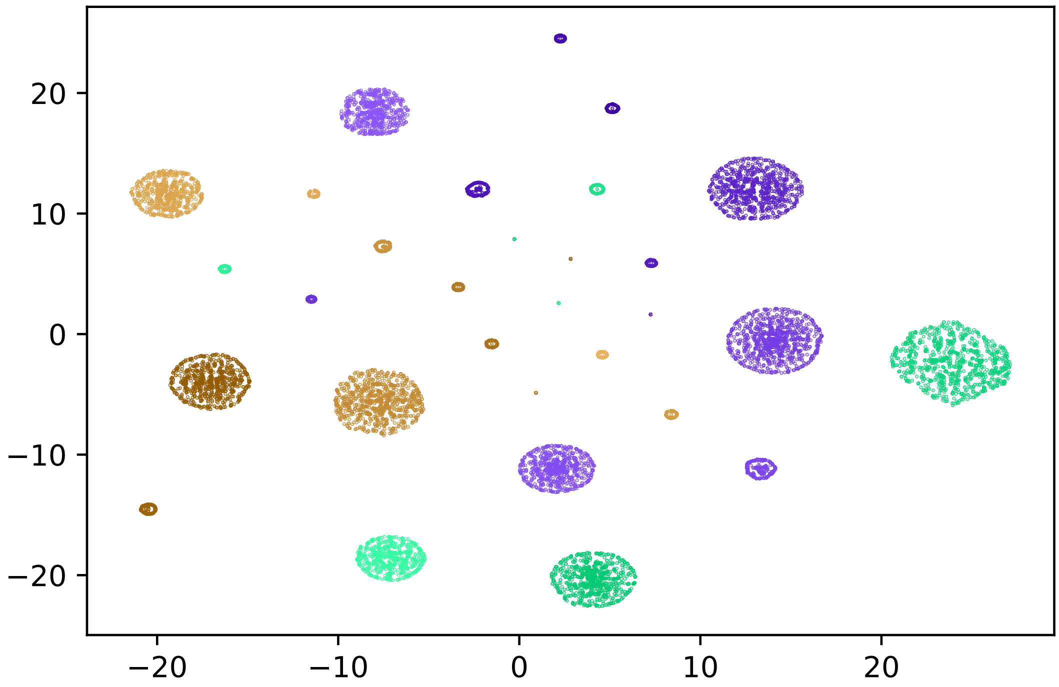


Figure 6: t-SNE plot for all the proteins (30) in the test set of DUD-E dataset. Each cluster is distinguishable with others and denotes a protein. Note that the SSnet model had no information about these proteins as they were in the test set and is yet able to distinguish them.

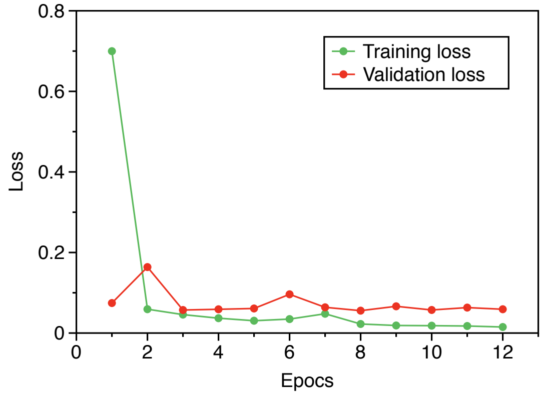


(a)

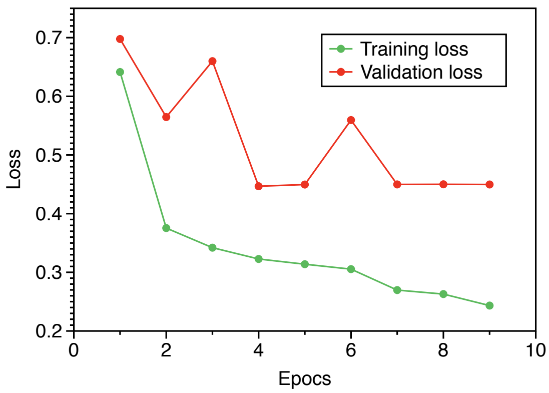


(b)

Figure 7: SSnet model training over a) DUD-E dataset and b) BindingDB dataset

### Hyper parameters in GNN-CNN

The parameters used for GNN-CNN are

- radius=2
- ngram=1
- dim=10
- layer_gnn=3
- side=5
- window=$((2*side+1))
- layer_cnn=3
- layer_output=3
- lr=1e-3
- lr_decay=0.5
- decay_interval=10
- weight_decay=1e-6
- iteration=100

Table 2: MUV data details

| MUV ID | PDB ID | Ligand | Ki/IC50 (nM) | Assay type |
|---|---|---|---|---|
| 600 | 1YOW | P0E | N/A | cell |
| 692 | 1YOW | P0E | N/A | cell |
| 859 | 5CXV | 0HK | N/A | cell |
| 852 | 4XE4 | NAG | N/A | biochemical |
| 548 | 3POO | S69 | N/A | biochemical |
| 832 | 1AU8 | 0H8 | N/A | biochemical |
| 689 | 2Y6O | 1N1 | 25 | biochemical |
| 846 | 5EXM | 5ST | N/A | biochemical |
| 466 | 3V2Y | ML5 | 18-77 | cell |

Table 3: $EF_{0.5\%}$ comparison on MUV dataset

| Target | 3D-CNN | vina | SSnet:DUD-E | GNN-CNN | SSnet:BDB |
|---|---|---|---|---|---|
| 692 | 0.000 | 0.000 | 0.000 | 0.000 | **6.680** |
| 859 | 0.000 | 0.000 | 0.000 | 0.000 | **13.360** |
| 548 | 0.000 | 0.000 | 0.000 | 0.000 | 0.000 |
| 600 | 0.000 | 0.000 | 0.000 | 0.000 | **6.680** |
| 852 | 0.000 | 0.000 | 0.000 | 0.000 | 0.000 |
| 846 | **6.667** | 0.000 | 0.000 | 0.000 | 0.000 |
| 832 | 0.000 | 0.000 | 0.000 | 0.000 | 0.000 |
| 689 | **6.667** | 0.000 | 0.000 | 0.000 | 0.000 |
| 466 | 0.000 | 0.000 | 0.000 | 0.000 | 0.000 |
| mean | 1.482 | 0.000 | 0.000 | 0.000 | **2.969** |
| std. dev. | 2.940 | 0.000 | 0.000 | 0.000 | 4.853 |

Table 4: $EF_{1.0\%}$ comparison on MUV dataset

| Target | 3D-CNN | vina | SSnet:DUD-E | GNN-CNN | SSnet:BDB |
|---|---|---|---|---|---|
| 692 | 0.000 | 0.000 | 0.000 | 0.000 | **3.340** |
| 859 | 0.000 | 0.000 | 0.000 | 0.000 | **6.680** |
| 548 | **6.667** | 0.000 | 3.318 | 0.000 | 0.000 |
| 600 | 0.000 | 0.000 | 0.000 | 0.000 | **3.340** |
| 852 | 0.000 | 0.000 | 0.000 | 0.000 | 0.000 |
| 846 | **3.330** | 0.000 | 0.000 | 0.000 | 0.000 |
| 832 | 0.000 | **3.330** | 0.000 | 0.000 | 0.000 |
| 689 | **3.330** | **3.330** | 0.000 | 0.000 | 0.000 |
| 466 | 0.000 | **3.330** | 0.000 | 0.000 | 0.000 |
| mean | **1.481** | 1.110 | 0.369 | 0.000 | **1.484** |
| std. dev. | 2.421 | 1.665 | 1.106 | 0.000 | 2.426 |

Table 5: $EF_{5.0\%}$ comparison on MUV dataset

| Target | 3D-CNN | vina | SSnet:DUD-E | GNN-CNN | SSnet:BDB |
|---|---|---|---|---|---|
| 692 | 0.000 | 0.000 | 0.000 | 0.000 | **2.001** |
| 859 | **2.000** | 1.330 | 1.332 | 1.665 | 1.334 |
| 548 | **6.667** | 0.000 | 2.665 | 0.690 | 0.667 |
| 600 | 1.330 | **2.667** | 1.999 | 0.667 | **2.668** |
| 852 | 0.000 | 0.667 | 0.666 | **1.378** | 0.000 |
| 846 | **0.667** | **0.667** | 0.000 | 0.000 | **0.667** |
| 832 | 0.000 | **3.330** | 0.000 | 1.332 | 0.667 |
| 689 | **2.000** | **2.000** | 1.332 | 1.379 | 0.000 |
| 466 | 1.330 | 1.330 | 0.666 | 0.000 | **2.001** |
| mean | **1.555** | 1.332 | 0.962 | 0.790 | 1.112 |
| std. dev. | 2.082 | 1.154 | 0.949 | 0.676 | 0.943 |

Table 6: AUC comparison on DUD-E test set

| Targets | Vina | 3D-CNN | GNN-CNN | | SSnet | |
|---|---|---|---|---|---|---|
| | | | DUD-E | BDB | DUD-E | BDB |
| ABL1 | 0.75 | 0.93 | 0.98 | 0.86 | 0.98 | 0.79 |
| ADRB1 | 0.74 | 0.88 | 0.98 | 0.78 | 0.99 | 0.71 |
| AKT1 | 0.74 | 0.98 | 0.99 | 0.91 | 1.00 | 0.91 |
| AKT2 | 0.78 | 0.99 | 0.98 | 0.94 | 0.99 | 0.93 |
| ANDR | 0.64 | 0.73 | 0.81 | 0.80 | 0.79 | 0.88 |
| BRAF | 0.84 | 0.99 | 0.98 | 0.91 | 0.99 | 0.86 |
| CDK2 | 0.72 | 0.84 | 0.93 | 0.86 | 0.94 | 0.85 |
| CP3A4 | 0.60 | 0.90 | 0.95 | 0.82 | 0.96 | 0.76 |
| DYR | 0.77 | 0.87 | 0.95 | 0.80 | 0.97 | 0.74 |
| FAK1 | 0.80 | 0.99 | 0.96 | 0.66 | 0.99 | 0.93 |
| FPPS | 0.29 | 0.98 | 0.99 | 0.96 | 0.99 | 0.70 |
| GRIA2 | 0.75 | 0.78 | 0.99 | 0.41 | 0.97 | 0.61 |
| HIVPR | 0.72 | 0.89 | 0.95 | 0.63 | 0.96 | 0.89 |
| ITAL | 0.60 | 0.94 | 0.96 | 0.93 | 0.87 | 0.91 |
| JAK2 | 0.77 | 0.99 | 0.94 | 0.86 | 0.99 | 0.94 |
| KPCB | 0.76 | 0.86 | 0.98 | 0.93 | 0.97 | 0.81 |
| LCK | 0.80 | 0.92 | 0.96 | 0.82 | 0.98 | 0.87 |
| LKHA4 | 0.82 | 0.94 | 0.96 | 0.88 | 0.99 | 0.74 |
| MK01 | 0.85 | 0.93 | 1.00 | 0.77 | 0.98 | 0.93 |
| NOS1 | 0.59 | 0.73 | 0.90 | 0.93 | 0.97 | 0.48 |
| PGH2 | 0.74 | 0.84 | 0.95 | 0.50 | 0.93 | 0.62 |
| PPARA | 0.87 | 0.87 | 0.94 | 0.51 | 1.00 | 0.89 |
| PPARG | 0.80 | 0.92 | 0.99 | 0.80 | 0.99 | 0.82 |
| PRGR | 0.68 | 0.85 | 0.98 | 0.79 | 0.90 | 0.94 |
| PYRD | 0.77 | 0.92 | 0.88 | 0.85 | 0.98 | 0.80 |
| SRC | 0.65 | 0.95 | 0.94 | 0.64 | 0.99 | 0.88 |
| THB | 0.75 | 0.83 | 0.98 | 0.88 | 0.96 | 0.83 |
| UROK | 0.77 | 0.96 | 0.98 | 0.79 | 0.99 | 0.63 |
| WEE1 | 0.83 | 0.99 | 0.98 | 0.66 | 1.00 | 0.95 |
| mean | 0.73 | 0.90 | 0.96 | 0.79 | 0.97 | 0.81 |
| std. dev. | 0.11 | 0.07 | 0.04 | 0.14 | 0.05 | 0.12 |

Table 7: $EF_{1\%}$ comparison on DUD-E test set

| Targets | Vina | 3D-CNN | GNN-CNN | | SSnet | |
|---|---|---|---|---|---|---|
| | | | DUD-E | BDB | DUD-E | BDB |
| ABL1 | 10 | 69 | 42 | 23 | 46 | 24 |
| ADRB1 | 5 | 19 | 36 | 14 | 50 | 7 |
| AKT1 | 8 | 85 | 48 | 29 | 53 | 33 |
| AKT2 | 15 | 67 | 33 | 31 | 42 | 36 |
| ANDR | 16 | 3 | 16 | 10 | 18 | 22 |
| BRAF | 18 | 78 | 42 | 25 | 56 | 18 |
| CDK2 | 6 | 26 | 25 | 21 | 32 | 18 |
| CP3A4 | 2 | 29 | 25 | 9 | 33 | 11 |
| DYR | 3 | 29 | 37 | 23 | 38 | 16 |
| FAK1 | 5 | 87 | 38 | 2 | 40 | 16 |
| FPPS | 0 | 46 | 35 | 23 | 59 | 4 |
| GRIA2 | 13 | 3 | 36 | 0 | 17 | 2 |
| HIVPR | 6 | 12 | 13 | 1 | 30 | 10 |
| ITAL | 0 | 41 | 26 | 27 | 19 | 26 |
| JAK2 | 10 | 86 | 26 | 5 | 58 | 26 |
| KPCB | 10 | 5 | 42 | 23 | 35 | 13 |
| LCK | 7 | 54 | 26 | 7 | 51 | 22 |
| LKHA4 | 7 | 36 | 44 | 17 | 45 | 7 |
| MK01 | 4 | 46 | 51 | 6 | 40 | 41 |
| NOS1 | 1 | 10 | 11 | 49 | 39 | 5 |
| PGH2 | 13 | 18 | 31 | 0 | 18 | 1 |
| PPARA | 9 | 12 | 17 | 0 | 47 | 13 |
| PPARG | 6 | 22 | 41 | 6 | 49 | 14 |
| PRGR | 9 | 7 | 42 | 10 | 27 | 30 |
| PYRD | 8 | 42 | 19 | 19 | 49 | 12 |
| SRC | 3 | 61 | 47 | 0 | 54 | 30 |
| THB | 12 | 9 | 53 | 27 | 34 | 15 |
| UROK | 9 | 54 | 46 | 3 | 45 | 4 |
| WEE1 | 8 | 77 | 39 | 4 | 56 | 19 |
| mean | 8 | 39 | 34 | 14 | 41 | 17 |
| std. dev. | 5 | 28 | 12 | 12 | 13 | 11 |

Table 8: AUC comparison on BDB test set

| PDB | SSnet:DUD-E | SSnet:BDB | GNN-CNN:DUD-E | GNN-CNN:BDB | Actives | Inactives | PLIs |
|---|---|---|---|---|---|---|---|
| 1CAH | 0.689 | 0.94 | 0.678 | 0.839 | 3688 | 1094 | 4782 |
| 1CQP | 0.334 | 0.90 | 0.251 | 0.818 | 258 | 145 | 403 |
| 1CVW | 0.634 | 0.91 | 0.564 | 0.799 | 397 | 146 | 543 |
| 1D3D | 0.712 | 0.91 | 0.617 | 0.855 | 2159 | 2702 | 4861 |
| 1D3G | 0.607 | 0.94 | 0.753 | 0.787 | 270 | 413 | 683 |
| 1D6O | 0.504 | 0.96 | 0.797 | 0.894 | 205 | 169 | 374 |
| 1DB4 | 0.773 | 0.88 | 0.800 | 0.889 | 114 | 145 | 259 |
| 1DHF | 0.503 | 0.87 | 0.578 | 0.893 | 402 | 288 | 690 |
| 1DI9 | 0.744 | 0.93 | 0.615 | 0.812 | 2447 | 448 | 2895 |
| 1DKF | 0.528 | 0.87 | 0.635 | 0.860 | 219 | 23 | 242 |
| 1E3G | 0.393 | 0.92 | 0.469 | 0.838 | 1670 | 275 | 1945 |
| 1E3K | 0.662 | 0.85 | 0.690 | 0.867 | 1365 | 39 | 1404 |
| 1ERE | 0.915 | 0.93 | 0.873 | 0.930 | 2116 | 767 | 2883 |
| 1EUB | 0.742 | 0.89 | 0.715 | 0.822 | 1715 | 230 | 1945 |
| 1EZQ | 0.550 | 0.93 | 0.530 | 0.807 | 3174 | 772 | 3946 |
| 1F9X | 0.848 | 0.83 | 0.723 | 0.911 | 393 | 512 | 905 |
| 1FAP | 0.645 | 0.93 | 0.412 | 0.913 | 2705 | 216 | 2921 |
| 1FBY | 0.635 | 0.87 | 0.479 | 0.904 | 667 | 42 | 709 |
| 1FKN | 0.499 | 0.92 | 0.499 | 0.785 | 4758 | 1175 | 5933 |
| 1FMK | 0.745 | 0.92 | 0.702 | 0.908 | 1401 | 1001 | 2402 |
| 1FT2 | 0.774 | 0.92 | 0.605 | 0.816 | 374 | 92 | 466 |
| 1GFW | 0.900 | 0.88 | 0.859 | 0.890 | 529 | 872 | 1401 |
| 1GZK | 0.518 | 0.89 | 0.367 | 0.846 | 495 | 49 | 544 |
| 1HRH | 0.648 | 0.90 | 0.594 | 0.829 | 428 | 738 | 1166 |
| 1HRN | 0.648 | 0.93 | 0.290 | 0.818 | 2728 | 239 | 2967 |
| 1HYV | 0.761 | 0.92 | 0.466 | 0.846 | 373 | 1547 | 1920 |
| 1I44 | 0.789 | 0.96 | 0.678 | 0.802 | 333 | 227 | 560 |
| 1I7G | 0.632 | 0.91 | 0.495 | 0.793 | 854 | 474 | 1328 |
| 1IAS | 0.697 | 0.92 | 0.428 | 0.767 | 919 | 88 | 1007 |
| 1IKV | 0.544 | 0.88 | 0.542 | 0.652 | 238 | 201 | 439 |
| 1JNK | 0.725 | 0.98 | 0.736 | 0.895 | 284 | 456 | 740 |
| 1KWP | 0.692 | 0.90 | 0.592 | 0.841 | 290 | 257 | 547 |
| 1NDE | 0.896 | 0.92 | 0.862 | 0.871 | 1147 | 607 | 1754 |
| 1O86 | 0.638 | 0.90 | 0.353 | 0.896 | 267 | 282 | 549 |
| 1S9I | 0.851 | 1.00 | 0.533 | 0.746 | 85 | 12 | 97 |
| 1S9J | 0.721 | 0.85 | 0.611 | 0.815 | 474 | 74 | 548 |
| 1T64 | 0.773 | 0.92 | 0.751 | 0.839 | 133 | 347 | 480 |
| 1UK0 | 0.814 | 0.93 | 0.740 | 0.909 | 2083 | 288 | 2371 |
| 1UWJ | 0.754 | 0.94 | 0.514 | 0.955 | 2176 | 150 | 2326 |
| 1V4S | 0.669 | 0.87 | 0.547 | 0.811 | 243 | 61 | 304 |
| 1W0E | 0.590 | 0.88 | 0.619 | 0.796 | 220 | 2557 | 2777 |
| 1Y6A | 0.574 | 0.91 | 0.500 | 0.863 | 3987 | 646 | 4633 |
| 2B7A | 0.323 | 0.91 | 0.453 | 0.959 | 5707 | 966 | 6673 |
| 2F2U | 0.757 | 0.93 | 0.590 | 0.875 | 1777 | 254 | 2031 |
| 2FZJ | 0.767 | 0.91 | 0.626 | 0.896 | 107 | 13 | 120 |
| 2KAV | 0.370 | 0.90 | 0.555 | 0.843 | 2307 | 723 | 3030 |
| 2M2F | 0.918 | 0.90 | 0.868 | 0.885 | 960 | 576 | 1536 |
| 3EML | 0.640 | 0.89 | 0.724 | 0.866 | 2214 | 392 | 2606 |
| 3MAX | 0.679 | 0.93 | 0.670 | 0.818 | 1707 | 666 | 2373 |
| 3O8Y | 0.630 | 0.96 | 0.588 | 0.830 | 198 | 417 | 615 |
| 3VG9 | 0.850 | 0.93 | 0.595 | 0.884 | 955 | 576 | 1531 |
| 4PH9 | 0.571 | 0.92 | 0.578 | 0.854 | 1184 | 1224 | 2408 |
| mean | 0.669 | 0.911 | 0.602 | 0.849 | 1267 | 513 | 1780 |
| std. dev. | 0.143 | 0.032 | 0.144 | 0.055 | 1292 | 557 | 1545 |

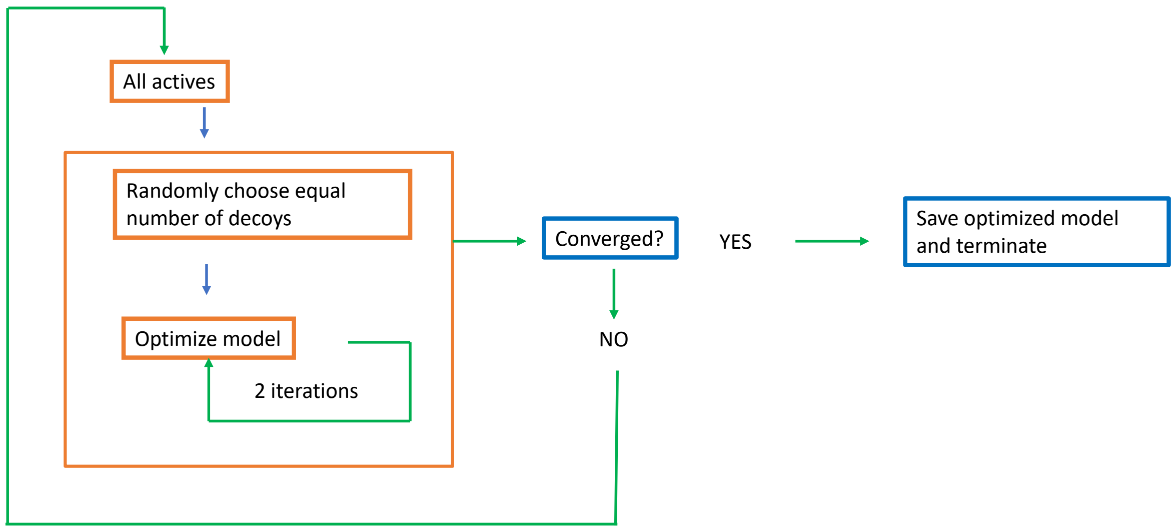


Figure 8: Dynamic model optimization for SSnet:DUD-E

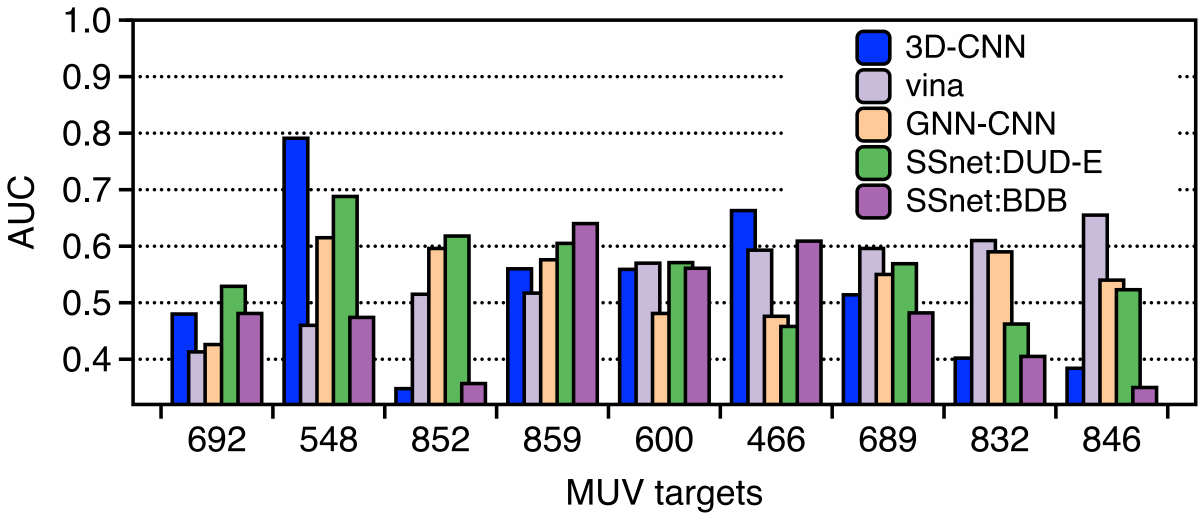


Figure 9: Various model performance on MUV targets for AUC

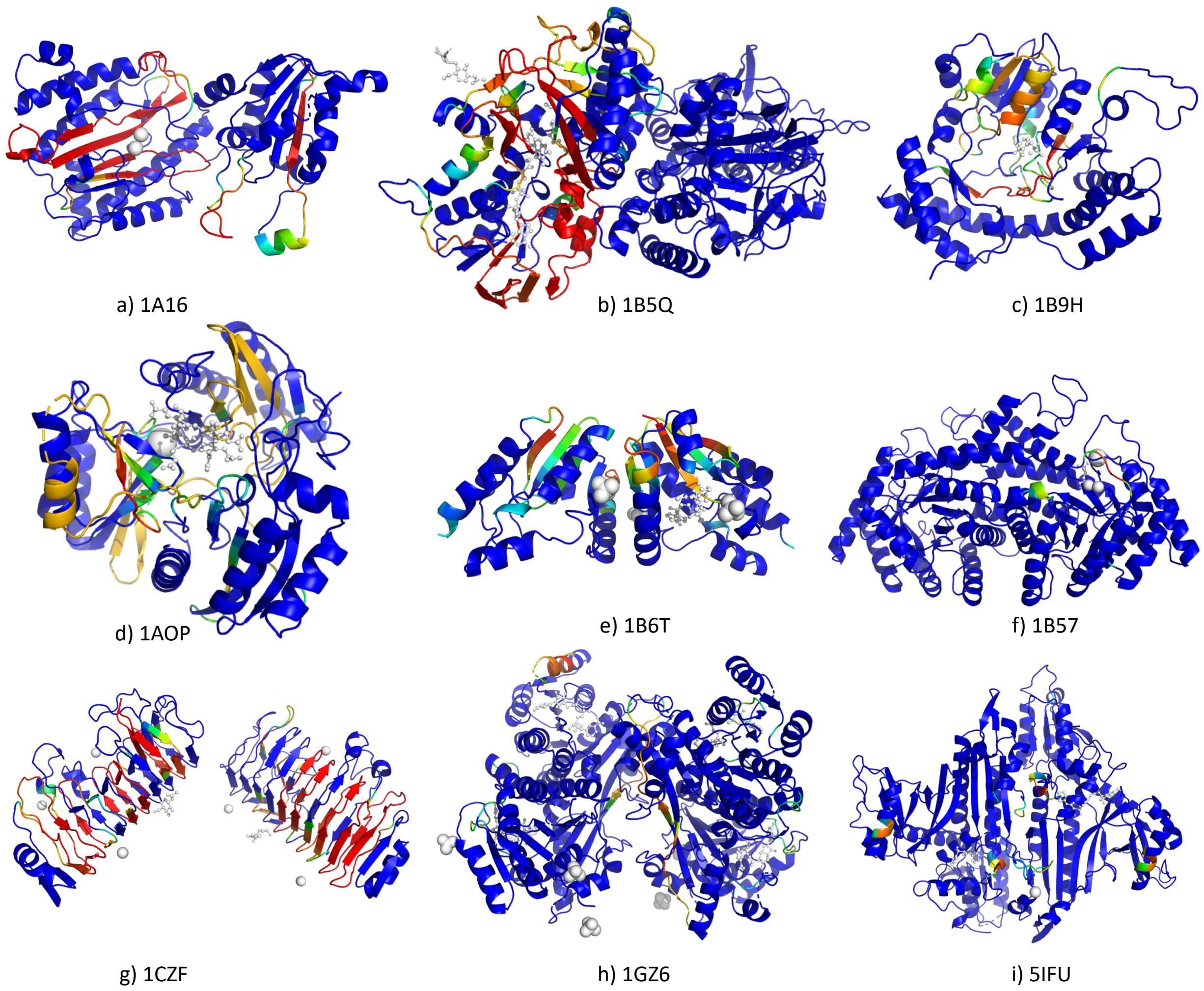


Figure 10: Grad-CAM visualization of the heatmap for nine different proteins with their PDB ID. The heatmap is a rainbow mapping with violet as the lowest and red as the highest value. The ligand and other small molecules are shown in grey.

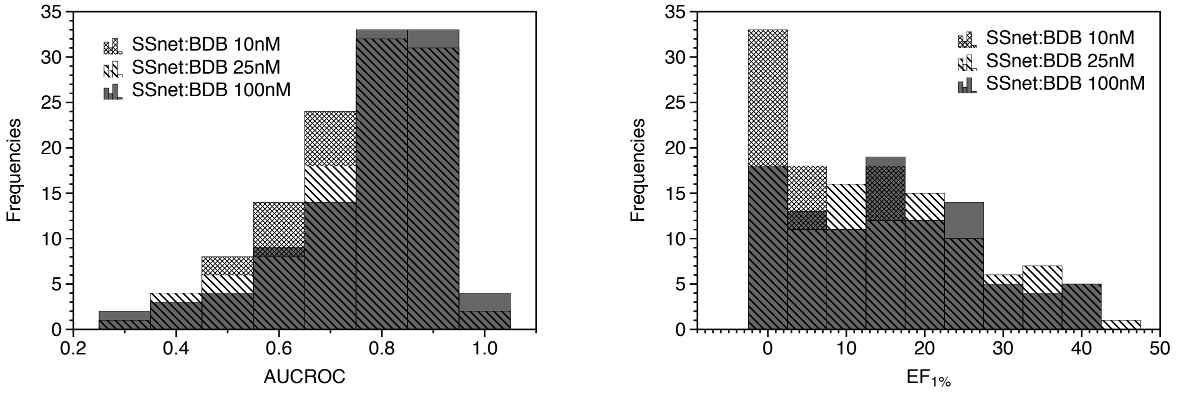


(a) (b)

Figure 11: SSnet:BDB performance when various cutoff for IC50 is applied and tested 102 targets of DUD-E dataset. The mean score for AUCROC are 0.77, 0.76 and 0.73 for 100, 25 and 10 nM cutoff respectively. The mean score for $EF_{1\%}$ are 15, 16 and 10 for 100, 25 and 10 nM cutoff respectively.
